# Impaired Macrophage DNase Secretion Promotes Neutrophil Extracellular Trap Mediated Defective Efferocytosis in Atherosclerosis

**DOI:** 10.1101/2025.01.30.635503

**Authors:** Umesh Kumar Dhawan, Tanwi Vartak, Hanna Englert, Aarushi Singhal, Rahul Chakraborty, Karran Kiran Bhagat, Luiz Ricardo C. Vasconcellos, Ciaran McDonnell, Mary Connolly, Edward Mulkern, Martin O’Donohoe, Mathias Gelderblom, Thomas Renne, Catherine Godson, Eoin Brennan, Manikandan Subramanian

## Abstract

Neutrophil extracellular traps (NETs) drive atherosclerosis progression and are associated with adverse clinical outcomes like myocardial infarction and stroke. While the triggers of NETosis in atherosclerotic plaque are well-characterized, the mechanisms underlying NET degradation and clearance remain unclear. Moreover, the impact of impaired NET clearance on atherosclerosis progression have not been elucidated. Here we show that macrophages are critical for the release of DNases, which degrade NETs. We identified endoplasmic reticulum (ER) stress mediated activation of PERK-ATF4 pathway as a key driver of impaired macrophage DNase secretion, leading to delayed NET clearance and their persistence. Elevated NET levels trigger cleavage of the efferocytosis receptor Mertk leading to defective macrophage efferocytosis and exacerbation of plaque necrosis. Human atherosclerotic plaques and *Ldlr^-/-^* mice treated with ISRIB, a PERK inhibitor, show enhanced DNase secretion and clearance of NETs. Together, the identification of key mechanisms of NET clearance in atherosclerosis offers new therapeutic strategies to stabilize plaques.

## Introduction

Atherosclerotic cardiovascular disease (CVD) remains a leading cause of morbidity and mortality worldwide^1,2^. Although lipid-lowering therapies have significantly reduced CVD-related deaths, a substantial residual risk for adverse clinical outcomes persists^3^. Defective efferocytosis, the process of phagocytic clearance of apoptotic cells^4^, and unresolved inflammation, are key drivers of the development of clinically dangerous necrotic atherosclerotic plaques^5^. Therefore, novel strategies to promote efferocytosis and resolve inflammation are of interest.

Recently, neutrophil extracellular traps (NETs), web-like DNA structures released by neutrophils during NETosis, have emerged as a key driver of inflammation in atherosclerotic lesions and contribute to plaque progression^6,7^. Several factors within the atherosclerotic plaque promote NETosis and release of NETs, including cholesterol crystals^8^, CCL7^9^, and lipid-induced oxidative stress^10^. Additionally, gene mutations associated with clonal hematopoiesis of indeterminate potential (CHIP) including LNK^11^, and JAK2^12–14^ increase the susceptibility of neutrophils to release NETs, thereby exacerbating plaque progression and increasing CVD mortality.

NETs are potent danger-associated molecular patterns (DAMPs) that elicit a robust pro-inflammatory response^15^. Therefore, the prompt degradation of NETs by extracellular endonucleases DNase1 and DNase1L3 is crucial for preventing tissue damage and resolving inflammation^16,17^. We previously identified a physiologic NET-induced DNase response pathway that operates in a feedback manner to maintain low NET levels in circulation. Moreover, we showed that NETs in systemic circulation trigger the release of DNases from liver and intestine to rapidly degrade NETs and restore homeostasis^18^. In contrast, the specific cells that express and release DNases in response to NETs generated within tissues, such as those in atherosclerotic plaques, are not characterized. Moreover, how the NET-induced DNase response is impaired upon plaque progression leading to accumulation of NETs is not understood. While higher levels of lesional NETs correlate directly with increased plaque necrosis and clinical complications, whether NETs directly contribute to efferocytosis impairment and necrotic core formation is not known.

In this study, we show that lesional macrophages are critical for the generation of DNases and degradation of NETs in the atherosclerotic plaque. Moreover, using a combination of human and mouse atherosclerosis, we demonstrate that lipid-induced endoplasmic reticulum (ER) stress in atherogenic macrophages triggers activation of the PERK-ATF4 signaling pathway to impair the NET-induced DNase response with consequent accumulation of NETs in advanced atherosclerosis. Finally, we demonstrate that NETs directly impair macrophage efferocytosis thereby contributing to necrotic core expansion and the formation of vulnerable plaques.

## Results

To characterize the lesional cells that express DNase1 and DNase1L3 and therefore could participate in clearance of NETs via secretion of DNases, we performed immunostainings for DNase1 and DNase1L3 along with cell type-specific markers for major lesional cell types including smooth muscle cells, macrophages, and endothelial cells. Murine atherosclerotic plaques showed expression of DNase1 and DNase1L3 in all lesional cell types examined (**Sup. Fig. 1A**). However, immunofluorescence signal identified macrophages as major source of plaque DNase (**Fig. 1A**). Indeed, both human and mouse macrophages release DNase1 and DNase1L3 in response to NETs^18^. Since the role of macrophage DNases in the clearance of NETs is not known, we compared the NET clearance efficiency of *Dnase1^-/-^ Dnase1l3^-/-^* (DKO) macrophages with WT macrophages. We observed that deficiency of DNase1 and DNase1L3 in macrophages impairs their ability to degrade NETs (**Fig. 1B**) as well as its subsequent engulfment by macrophages (**Fig. 1C**). Based on these data, we hypothesized that macrophages are critical for clearance of NETs in atherosclerotic plaques via the release of DNases.

**Fig. 1.**
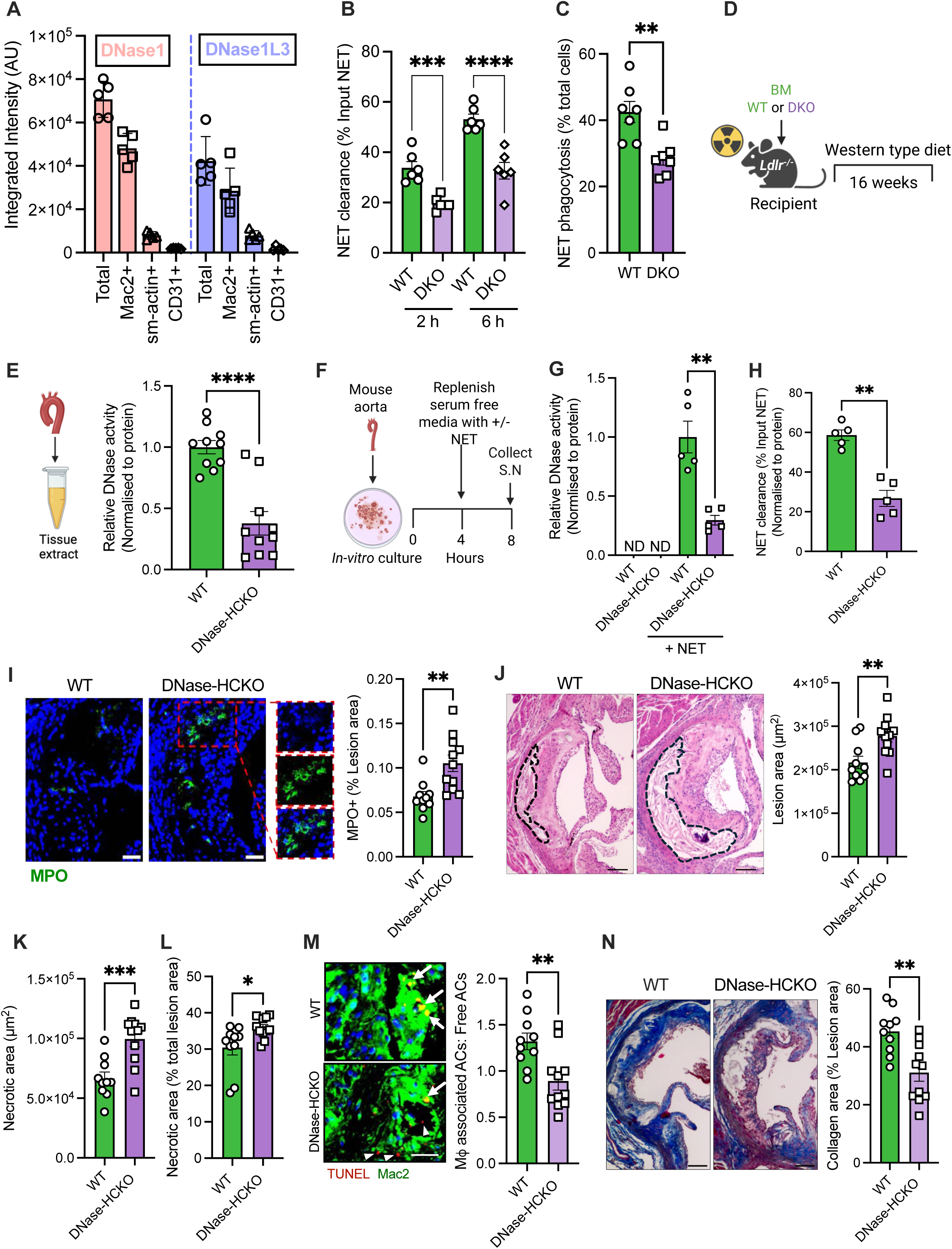
DNase deficiency in hematopoietic cells impairs NET clearance in atherosclerotic plaques. **(A)** The relative levels of DNase1 and DNase1L3 in lesional macrophages (Mac2+), smooth muscle cells (sm-actin +), and endothelial cells (CD31+) was quantified by measuring DNase1 and DNase1L3 fluorescence intensity in antibody-labelled aortic root sections of 16-week WD-fed *Ldlr^-/-^* mice. **(B)** BMDMs from WT and *Dnase1^-/-^Dnase1l3^-/-^* (DKO) mice were exposed to NETs (250 ng/ml) for either 2 h or 6 h. The quantity of remaining NETs in the supernatant was measured by Picogreen assay and the NET clearance efficiency was determined relative to input NET concentration. **(C)** BMDMs from WT and DKO mice were incubated for 2 hours with fluorescently labelled NETs. After washes, fluorescence microscopy was performed and the percent macrophages that showed engulfment of fluorescent NETs was quantified. **(D)** 8-week-old female *Ldlr^-/-^* mice were lethally irradiated followed by administration of bone marrow cells from either WT or DKO mice. Six weeks post bone marrow reconstitution, WT and DNase-hematopoietic cell knockout (DNase-HCKO) mice were fed a western-type diet for 16 weeks and then euthanized for analysis. **(E)** Brachiocephalic artery from WT and DNase-HCKO mice were homogenized, and the tissue extract was analyzed for DNase activity by SRED assay. n = 10 mice per group. **(F)** Aortic tree from both groups of mice (n = 5) were harvested, fragmented, and maintained in culture as explants followed by exposure to vehicle or NETs for 4 hours. The supernatant was collected for measurement of **(G)** DNase activity and **(H)** NET clearance efficiency. **(I)** Lesional NET level was quantified by immunostaining for myeloperoxidase (MPO) in DAPI-stained aortic root sections of WT and DNase-HCKO mice. n = 10 mice per group. Scale bar, 50 µm. **(J)** H&E-stained aortic root sections were analyzed for total atherosclerotic lesion area, **(K)** necrotic area, and **(L)** plaque necrosis as a percentage of total lesion area. The regions of plaque necrosis are demarcated by the black dashed line. n = 10-11 mice per group. Scale bar, 50 µm. **(M)** In-situ efferocytosis assay in aortic root sections labelled with TUNEL reagent to detect apoptotic cells (red) followed by Mac2 immunolabelling (green) to identify lesional macrophages. Lesional efferocytosis efficiency was calculated as the ratio of macrophage-associated apoptotic cells to free-lying apoptotic cells. White arrows indicate apoptotic cells associated with macrophages while white arrowheads show free-lying TUNEL+ cells. n = 10 mice per group. Scale bar, 25 µm. **(N)** Quantification of lesional collagen content in Mason’s trichrome-stained aortic root sections. n = 10 mice per group. Scale bar, 50 µm. All data are represented as mean ± SEM. Test for normality was conducted by Shapiro-Wilk test. P value was determined ANOVA (B), Mann-Whitney U test (C, G, and H), or unpaired t-test (D, I-N). *, p < 0.05; **, p < 0.01.

To test this hypothesis, we generated bone marrow chimeric mice that are deficient in DNase1 and DNase1L3 in all cells of hematopoietic origin by adoptively transferring bone marrow cells obtained from *Dnase1^-/-^Dnase1l3^-/-^* mice into lethally irradiated atherosclerosis prone *Ldlr^-/-^* mice (DNase-HCKO). Control group were injected bone marrow cells obtained from WT mice. Six weeks after bone marrow transplantation, both groups of mice were fed a western type diet for 16 weeks to generate advanced atherosclerosis (**Fig. 1D**). Aortic extracts were analyzed for DNase activity by single radial enzyme diffusion (SRED) assay.

Interestingly, we observed that the basal vascular DNase activity was lower in the DNase-HCKO mice as compared with WT mice (**Fig. 1E**) suggesting that hematopoietic cell derived DNase is critical for maintenance of basal tissue DNase activity. Important to note that plasma level of NETs (**Sup. Fig. 1B**) as well as plasma DNase activity was unaffected in the chimeric mice (**Sup. Fig. 1C**) demonstrating that hematopoietic cell derived DNases do not contribute significantly towards the maintenance of circulatory DNase activity.

Next, to address the question of relative contribution of hematopoietic vs. non-hematopoietic cells in the release of DNase in response to NETs within the atherosclerotic artery, we developed an *ex vivo* assay wherein atherosclerotic vascular tissue explants obtained from 16-week WD-fed WT or DNase-HCKO mice were exposed to either NETs or vehicle and the extent of release of DNase in the supernatant was quantified (**Fig. 1F**). In the absence of exogenously added NETs, the explant tissue did not release detectable levels of DNase (**Fig. 1G**). However, when NETs were added to the explant culture, DNase was released into the supernatant demonstrating the presence of an active NET-induced DNase response in the vascular explant tissue (**Fig. 1G**). Interestingly, compared with the WT mice, there was an ∼ 70% decrease in DNase activity in the supernatant of aortic tissue obtained from the DNase-HCKO mice (**Fig. 1G**). More importantly, this decrease in DNase activity was associated with an impairment in clearance of exogenously added NETs (**Fig. 1H**).

Based on these data, we questioned whether the decrease in plaque DNase activity in the DNase-HCKO mice is associated with increased accumulation of NETs. Indeed, DNase-HCKO mice showed increased signal of lesional MPO-DNA complexes in immunofluorescence indicative of increased accumulation of NETs (**Fig. 1I**). This accumulation is primarily driven by impaired DNase-mediated NET clearance since both WT and DKO neutrophils are equally susceptible to undergo NETosis when exposed to classical NETosis inducers such as PMA (**Sup. Fig. 1D**). From a disease perspective, the impaired clearance of NETs in the DNase-HCKO mice was associated with an increase in atherosclerotic plaque area (**Fig. 1J**). The increase in lesion area was not due to metabolic perturbations since body weight, plasma cholesterol, triglycerides, and blood glucose were similar between the WT and DNase-HCKO mice (**Sup. Fig. 1E-H**). More importantly, plaque necrosis, which is a critical determinant of vulnerable plaque formation, was increased in the DNase-HCKO mice (**Fig. 1K**). This increase was not due to larger lesions in the DNase-HCKO mice since the percentage lesion area that is necrotic was also higher (**Fig. 1L**). Since plaque necrosis is driven primarily by a defective macrophage efferocytosis^19^, we tested whether lesional efferocytosis efficiency is impaired in the DNase-HCKO mice using an *in-situ* efferocytosis assay^20^. Accordingly, aortic root sections were labeled with TUNEL reagent to detect intimal apoptotic cells and then co-stained with Mac2 to detect macrophages. The ratio of TUNEL+ nuclei that are associated with a macrophage to those that are lying free was used as an indicator of the *in-situ* efferocytosis efficiency^21^. Consistent with the plaque necrosis data, the DNase-HCKO mice demonstrated significant impairment in macrophage efferocytosis efficiency (**Fig. 1M**). Moreover, the DNase-HCKO mice showed decreased lesional collagen content (**Fig. 1N**). Taken together, these data demonstrate that DNase-HCKO mice have an accelerated atherosclerotic plaque progression with characteristics of plaque instability such as decreased collagen and larger necrotic area which in humans is associated with adverse clinical outcomes such as myocardial infarction and stroke.

### DNase deficiency leads to NET accumulation and impairment of macrophage efferocytosis

Since the accumulation of NETs in the DNase-HCKO mice was associated with decreased lesional efferocytosis efficiency, we asked whether NETs directly impair macrophage efferocytosis. Indeed, when macrophages were incubated with NETs *in vitro*, there was a concentration-dependent decrease in the uptake of apoptotic cells (**Fig. 2A**) suggesting that NETs decrease the efficiency of apoptotic cell clearance. Consistently, the NET-induced decrease in efferocytosis was more pronounced in the *Dnase1^-/-^Dnase1l3^-/-^* (DKO) macrophages (**Fig. 2B and Sup. Fig. 2A**) which have impaired NET clearance (**Fig. 1B**) and therefore persistently higher concentrations of NETs. WT and DKO macrophages have similar efferocytosis efficiency in the absence of NETs (**Sup. Fig. 2B**) demonstrating that the DKO macrophages do not have an intrinsic defect in efferocytosis. Taken together, these data suggest that a decrease or loss of NET-induced DNase secretion by macrophages leads to decrease DNase-mediated clearance and accumulation of NETs which impairs macrophage efferocytosis.

**Fig. 2.**
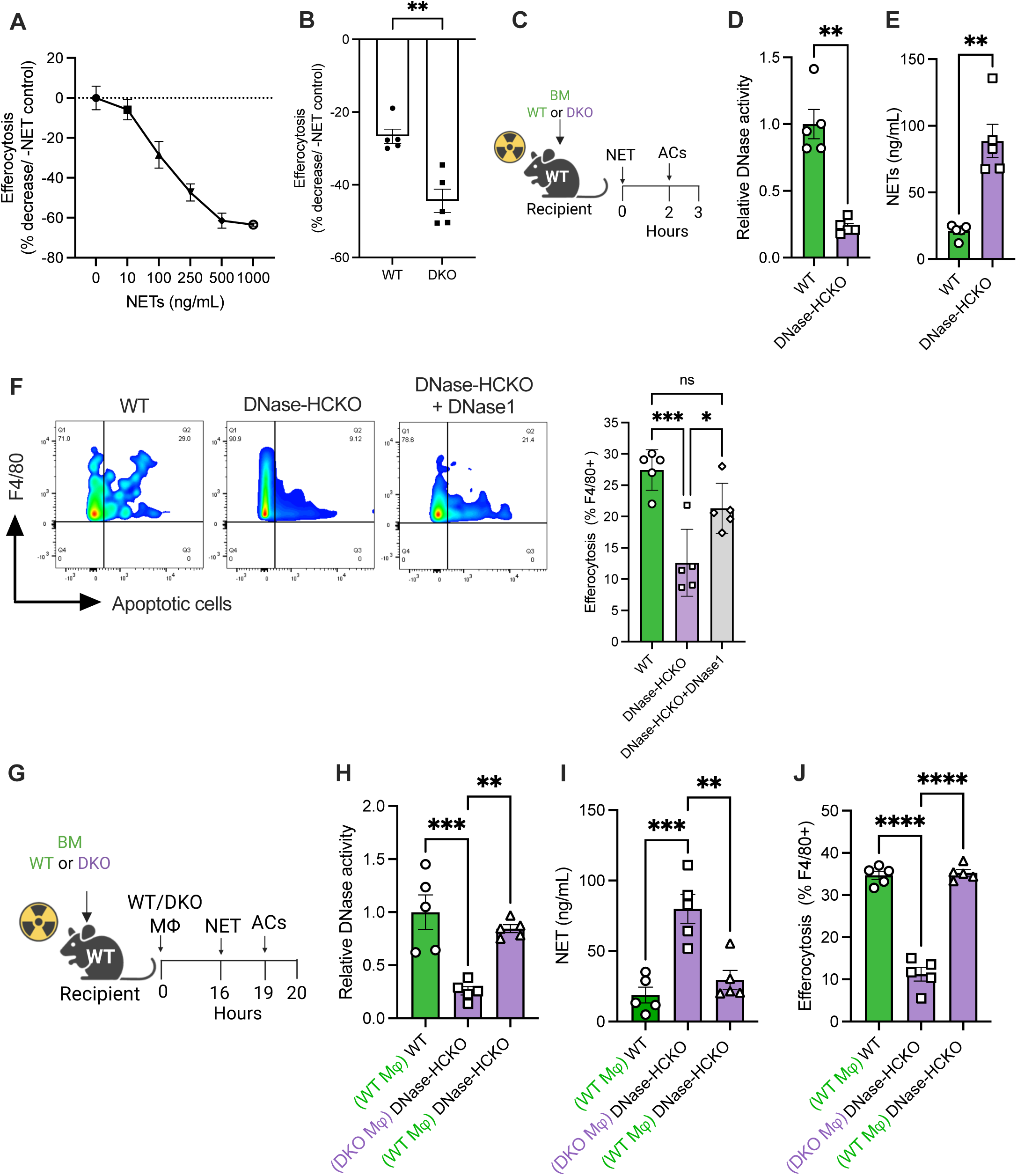
Impaired NET clearance promotes defective macrophage efferocytosis. **(A)** BMDMs were treated with vehicle or NETs at indicated concentrations for 2 h followed by incubation with fluorescently labelled apoptotic cells for 1 h. Efferocytosis efficiency was quantified by fluorescence microscopy and expressed as a percentage decrease relative to the vehicle group. n = 4 biological replicates. **(B)** WT and DKO BMDMs were exposed to vehicle or NETs (250 ng/ml) for 2 hours, then incubated with apoptotic cells for 1 hour. Efferocytosis efficiency is shown as a percentage change relative to the vehicle group. n = 5 biological replicates. **(C)** 8-week-old female C57BL/6J mice were lethally irradiated followed by injection of bone marrow cells from either WT or DKO mice. Six weeks post-transplant, WT and DNase-HCKO mice were injected intraperitoneally with NETs. A subgroup of DNase-HCKO mice also received purified DNase1 with NETs. Two hours later, CellVue Claret-labeled apoptotic cells were injected intraperitoneally. Mice were euthanized 1 hour later and peritoneal lavage was analyzed for **(D)** DNase activity by SRED, **(E)** NET levels by picogreen assay, and **(F)** macrophage efferocytosis efficiency by flow cytometry. N = 5 mice per group. **(G)** WT and DNase-HCKO chimeric mice were generated as above. WT mice received adoptive transfer of WT macrophages, while DNase-HCKO mice received WT or DKO macrophages. 16 h post adoptive cell transfer, the mice were injected NETs followed by apoptotic cells as described above for the measurement of peritoneal lavage **(H)** DNase activity, **(I)** NET levels, and **(J)** efferocytosis efficiency. n = 5 mice per group. The data are represented as mean ± SEM. Data were tested for normal distribution using Shapiro-Wilk test. P values were calculated using Mann Whitney U test (B, D, and E) and ANOVA with Tukey’s multiple comparisons test (F, H-J). *, p < 0.05; **, p < 0.01; ***, p < 0.001.

Next, to address the question whether accumulation of NETs impairs macrophage efferocytosis *in vivo*, we injected NETs into the peritoneal cavity of bone marrow chimeric C57BL/6 mice that were reconstituted with either WT or DKO bone marrow cells. Two hours later, the mice were given intraperitoneal injection of fluorescently labelled apoptotic cells.

One hour after AC injection, peritoneal lavage was performed (**Fig. 2C**). As with the atherosclerotic mice, the DNase-HCKO mice had decreased levels of DNase in the peritoneal lavage as compared with the WT mice (**Fig. 2D**). Most importantly, the decreased DNase activity in the DNase-HCKO mouse was associated with higher levels of uncleared NETs (**Fig. 2E**) and decreased macrophage efferocytosis (**Fig. 2F**), which could be rescued by administration of intraperitoneal DNase1 (**Fig. 2F**). These data suggest that loss of NET-induced production of DNases by hematopoietic cells in tissues leads to delayed clearance and accumulation of NETs with consequent impairment in macrophage efferocytosis.

Finally, to directly test whether macrophages are the major cellular source of the NET-induced DNase response in tissues, we adoptively transferred equal numbers of either WT or DNase deficient macrophages into the peritoneal cavity of DNase-HCKO mice (**Fig. 2G**). To ensure that the number of macrophages injected are within the physiological range, the mice were administered 0.5×10^6^ cells which represent half the number of resident peritoneal cavity macrophages. Consistent with our hypothesis, DNase-HCKO mice that received WT macrophages, but not DNase-deficient macrophages, had increased DNase activity which was comparable with the WT mice (**Fig. 2H**). Most importantly, these mice efficiently degraded the injected NETs (**Fig. 2I**) and was associated with improved macrophage efferocytosis (**Fig. 2J**). Overall, these data suggest that macrophages are critical for the NET-induced DNase response and play a dominant role in the clearance of locally generated NETs within the inflamed tissues.

### NETs impair macrophage efferocytosis via cleavage of the efferocytic receptor Mertk

Since efferocytosis is a multi-step process involving an initial step of recognition and binding of apoptotic cells followed by a second step of internalization^22^, we tested which of these broad processes are affected in the presence of NETs. To specifically test the efficiency of apoptotic cell binding, macrophages exposed to either vehicle or NETs were incubated with cytochalasin D, an actin polymerization inhibitor that blocks engulfment^23^, prior to addition of apoptotic cells. Interestingly, macrophages exposed to NETs were less efficient at binding apoptotic cells (**Fig. 3A**). Since efferocytosis receptors are critical for recognition and binding of ACs, we profiled the cell surface expression levels of major efferocytosis receptors including Mertk, Tim4, Axl, and LRP-1 by flow cytometry. While the levels of Tim4, Axl, and LRP-1 were unaffected (**Sup. Fig. 2C**), macrophages exposed to NETs demonstrated decreased cell surface expression of Mertk (**Fig. 3B**), a dominant apoptotic cell recognition receptor in macrophages in several tissues^24^. Cell surface expression of Mertk is regulated by post-translational modifications including proteolytic cleavage of its ectodomain resulting in the release of “soluble-Mertk” which is associated with loss of receptor function^25^. Indeed, the levels of soluble-Mertk was higher in the cell culture supernatants of macrophages incubated with NETs and showed a clear dose response (**Fig. 3C**). Consistent with these findings, the accumulation of NETs when incubated with DNase-deficient macrophages (**Fig. 1B**) was associated with increased generation of soluble-Mertk (**Fig. 3D**). Further supporting these *in vitro* data, the increased accumulation of NETs in the atherosclerotic plaques of DNase-HCKO mice (**Fig. 1I**) was associated with decreased expression of Mertk in Mac2+ lesional macrophages (**Fig. 3E**). Similarly, the increased persistence of NETs in the peritoneal cavity of DNase-HCKO mice (**Fig. 2E**) was associated with increased soluble-Mertk levels (**Fig. 3F**), and this effect could be reversed upon promoting NET degradation via administration of either DNase1 (**Fig. 3F**) or the adoptive transfer of WT macrophages (**Fig. 3G**). More importantly, the blockage of soluble-Mertk production and the preservation of cell surface Mertk levels in all these experimental conditions prevented the NET-induced impairment of macrophage efferocytosis (**Fig. 2F and 2G**).

**Fig. 3.**
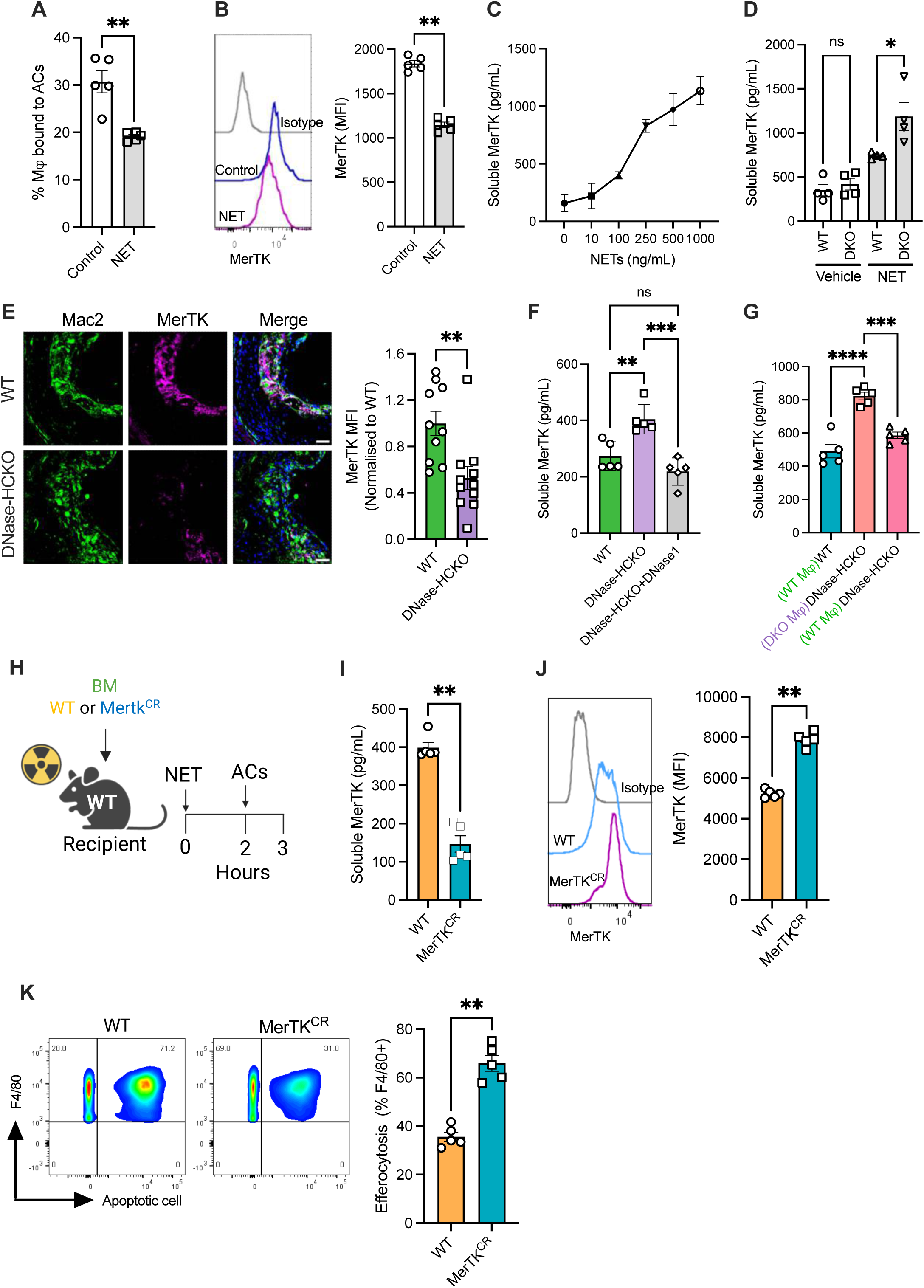
NETs cleave Mertk to impair efferocytosis. **(A)** BMDMs were exposed to NETs (250 ng/ml), then incubated with cytochalasin D (1 µM) for 30 min, followed by addition of fluorescently labelled apoptotic cells. After washes, apoptotic cell binding efficiency was quantified by fluorescence microscopy. n = 5 biological replicates. **(B)** Quantification of cell surface MerTK levels in BMDMs treated with or without NETs for 2 hours. n = 5 biological replicates. **(C)** ELISA-based measurement of soluble-Mertk in the cell culture supernatants of BMDMs exposed for 2 h with indicated concentrations of NETs. n = 3 biological replicates. **(D)** Quantification of soluble-Mertk in the cell culture supernatant of WT and DKO BMDMs exposed to vehicle or NETs (250 ng/ml) for 2 h. n = 4 biological replicates. **(E)** Quantification of Mertk levels in lesional macrophages (Mac2+) by immunostaining of aortic root sections of 16 wk-WD fed bone marrow chimeric WT and DNase-HCKO *Ldlr^-/-^* mice. n = 10 mice per group. **(F)** ELISA-based analysis of soluble-Mertk levels in the peritoneal lavage of bone marrow chimeric WT and DNase-HCKO mice two hours after intraperitoneal injection of NETs without or with DNase1. n = 5 mice per group. **(G)** As above, except that WT and DNase-HCKO mice were adoptively transferred either WT or DKO macrophages intraperitoneally 16 h prior to injection of NETs. n = 5 mice per group. **(H)** 8-week-old female *C57BL/6J* mice were lethally irradiated and injected bone marrow cells from either WT or Mertk cleavage resistant (Mertk^CR^) mice. Six weeks later, bone marrow chimeric mice were injected NETs (1 µg i.p.), and fluorescently labelled apoptotic cells 2 h later. Peritoneal lavage was performed 1 h later for analysis of **(I)** soluble-Mertk by ELISA; **(J)** macrophage Mertk expression by flow cytometry, and **(K)** macrophage efferocytosis efficiency by flow cytometry. n = 5 mice per group. The data are represented as mean ± SEM. Data were tested for normal distribution using Shapiro-Wilk test. P values were calculated using Mann Whitney U test (A, B, E, I-K) and ANOVA with Tukey’s multiple comparisons test (C, F, G). *, p < 0.05; **, p < 0.01; ***, p < 0.001.

We previously reported the identification of the cleavage site residues of Mertk^25^ which were mutated to generate a Mertk cleavage-resistant (Mertk^CR^) knockin mouse that was resistant to ADAM17-mediated proteolysis and shedding^26^. We used these Mertk cleavage-resistant mice to address the question whether NET-induced Mertk cleavage is a causal mechanism driving defective macrophage efferocytosis. In this context, we generated bone marrow chimeric C57BL/6 mice that were reconstituted with cells from either WT or Mertk^CR^ mouse. Both groups of mice were injected with NETs followed by fluorescently labeled apoptotic cells (**Fig. 3H**). As expected, the Soluble Mertk levels in the peritoneal lavage of the Mertk^CR^ mouse was significantly lower compared with the WT group (**Fig. 3I**). Consistent with the decreased generation of Soluble Mertk, the Mertk^CR^ group showed increased cell surface Mertk levels in the F4/80^+^ peritoneal cavity macrophages (**Fig. 3J**). Most importantly, the efferocytosis efficiency of Mertk^CR^ mice was significantly higher than the WT group (**Fig. 3K**). Taken together, these data demonstrate that cleavage of Mertk and the loss of this key apoptotic cell recognition receptor drives the NET-induced impairment of macrophage efferocytosis.

### NET-associated HMGB1 signals Mertk cleavage

We next explored the molecular mechanisms by which NETs trigger Mertk cleavage. First, we tested whether NETs activate ADAM17 to induce Mertk cleavage. Indeed, when macrophages were exposed to NETs in the presence of TAPI-0, an ADAM17 inhibitor^25^, both the generation of Soluble Mertk and the decrease in cell surface Mertk was abrogated (**Fig. 4A-B**), along with preservation of macrophage efferocytosis efficiency (**Fig. 4C**). These data demonstrate that NETs promote cleavage and shedding of Mertk via activation of the classical ADAM17 pathway.

**Fig. 4.**
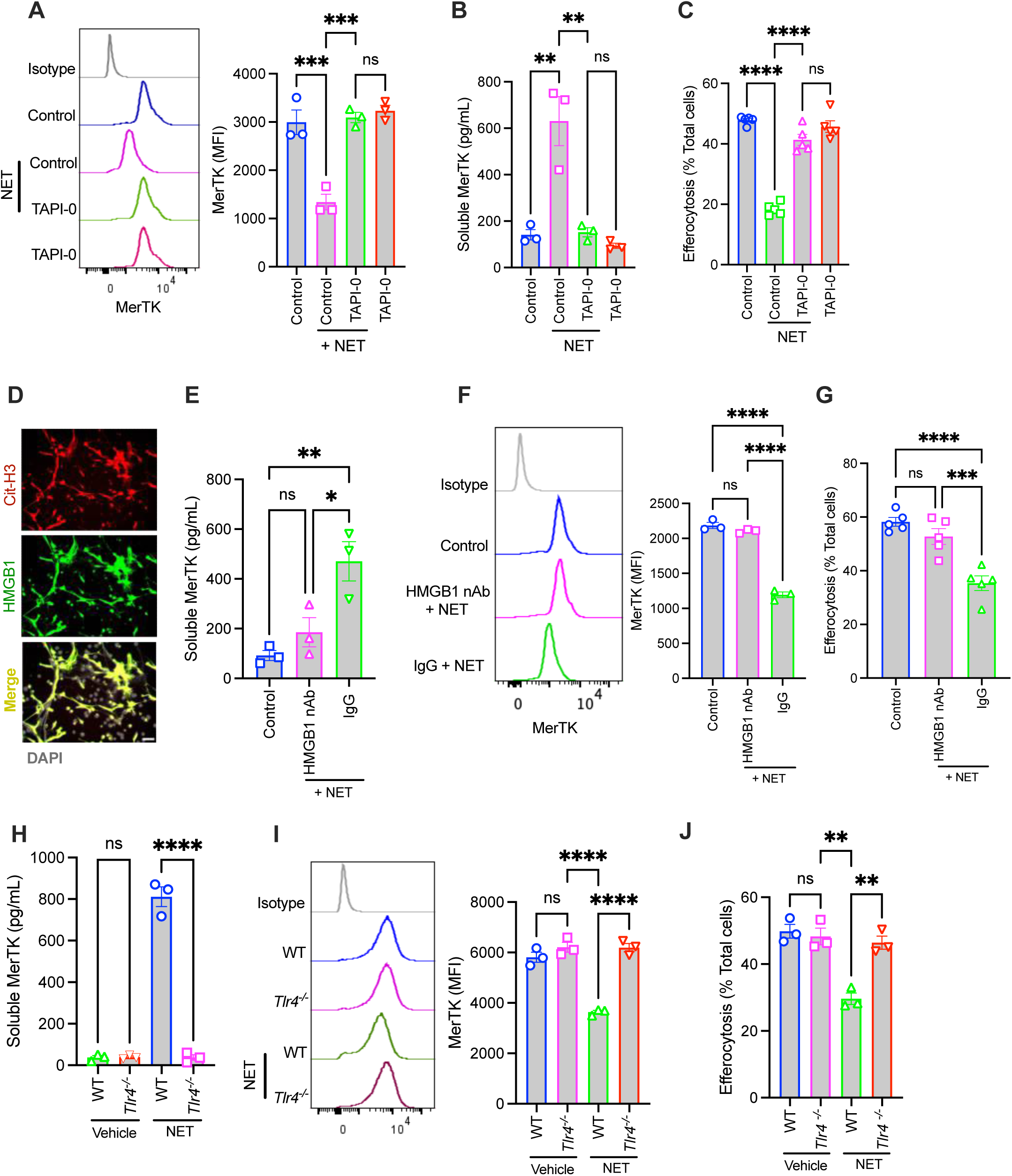
NET-associated HMGB1 triggers Mertk cleavage by TLR4-mediated activation of ADAM17. **(A)** BMDMs were pre-treated with TAPI-0 (10 µM) or vehicle for 1 h and then exposed to NETs for 2 h. The cell surface Mertk levels was quantified by flow cytometry. n = 3 biological replicates. **(B)** As above, except that the culture supernatant was assayed by ELISA for soluble-Mertk. n = 3 biological replicates. **(C)** Similar to (A), except that TAPI-0 and NET-exposed BMDMs were incubated with fluorescently labeled apoptotic cells was quantification of efferocytosis efficiency by microscopy. n = 5 biological replicates. **(D)** *In-vitro* generated NETs were immunostained with anti-CitH3 and anti-HMGB1 antibody and imaged by fluorescence microscopy. Nuclei were stained with DAPI. Scale bar, 50 µm. **(E)** BMDMs were exposed to NETs in the presence of a HMGB1 neutralizing Ab or control IgG. Soluble-Mertk was measured in the culture supernatants. n = 3 biological replicates. **(F)** Similar to (E), except that cell surface Mertk expression was quantified by flow cytometry. N = 3 biological replicates. **(G)** Efferocytosis efficiency was measured in BMDMs exposed to NETs in the presence of control IgG or HMGB1 neutralizing Ab. n = 5 biological replicates. **(H)** WT or *Tlr4^-/-^* BMDMs were exposed to NETs followed by quantification of soluble-Mertk levels in the supernatant, and **(I)** Mertk expression on the cell surface. n = 3 biological replicates. **(J)** Quantification of efferocytosis in WT and *Tlr4^-/-^* BMDMs exposed to vehicle or NETs. n = 3 biological replicates. The data are represented as mean ± SEM. P values were calculated using ANOVA with Tukey’s multiple comparisons test (A-C, E-J). *, p < 0.05; **, p < 0.01; ***, p < 0.001.

We then examined how NETs activate macrophage ADAM17. In this context, mass spectrometry analysis revealed HMGB1 as one of the dominant NET-associated proteins (**Sup. Fig. 3A**). This was further confirmed by both immunostaining (**Fig. 4D**) and western blotting (**Sup. Fig. 3B**) which demonstrated the presence of HMGB1 on NETs. These findings were particularly intriguing since HMGB1 is a damage-associated molecular pattern known to activate TLR4^27^ which is upstream of the TRIF-ROS-p38 pathway that leads to ADAM17 activation^25^. Indeed, incubation of macrophages with recombinant HMGB1 led to Mertk cleavage and decreased efferocytosis efficiency (**Sup. Fig. 3C-D**). We therefore tested whether NET-associated HMGB1 is relevant for Mertk cleavage. Interestingly, the incubation of macrophages with NETs in the presence of a HMGB1 neutralizing antibody blocked the production of NET-induced Soluble Mertk (**Fig. 4E**), enhanced cell surface Mertk (**Fig. 4F**), and restored macrophage efferocytosis efficiency (**Fig. 4G**). Consistent with these data,

HMGB1-deficient NETs, derived from HL60 cells subjected to siRNA mediated knockdown of HMGB1 (**Sup. Fig. 3E**), were unable to induce Mertk cleavage (**Sup. Fig. 3F**) or impair efferocytosis in macrophages (**Sup. Fig. 3G**). These data established HMGB1 as a NET-associated protein triggering Mertk cleavage. Finally, we tested whether HMGB1 signals via TLR4, a known activator of ADAM17, to promote Mertk cleavage. Indeed, macrophages from *Tlr4^-/-^* mice were resistant to HMGB1-induced generation of Soluble Mertk, loss of cell surface Mertk, and impairment of efferocytosis (**Sup. Fig. 3H-J**). More importantly, the lack of TLR4 in macrophages abrogated the ability of NETs to induce Mertk cleavage (**Fig. 4H-I**) and block efferocytosis (**Fig. 4J**). These data demonstrate that NET-associated HMGB1 is critical for TLR4-mediated activation of Mertk cleavage and impairment of macrophage efferocytosis.

### ATF4 activation in ER stressed atherogenic macrophages impairs their NET-induced DNase response

We showed previously that macrophages exposed to 7-ketocholesterol (7-KC), a modified form of cholesterol that is enriched in atherosclerotic plaques, induces ER stress leading to impairment of NET-induced DNase^18^. Importantly, we now show that the increased accumulation of NETs (**Fig. 5A-B**) in this pathophysiologically relevant state mimicking macrophages in an atherogenic environment is associated with increased Mertk cleavage (**Fig. 5C**) and defective efferocytosis (**Fig. 5D**), both of which could be reversed by ER stress relieving agents such as TUDCA and PBA (**Fig. 5C-D**). To understand the specific ER stress signaling cascade that mediates the impairment in the NET-induced DNase response, 7-KC-exposed macrophages were treated with either ISRIB, 4µ8C, or CeapinA7, specific inhibitors of the PERK, IRE1α, and ATF6 branch of the ER stress pathway, respectively, and the DNase activity in the supernatant was measured. Interestingly, the 7-KC-induced impairment in NET clearance was reversed specifically in macrophages exposed to the PERK inhibitor ISRIB, but not when exposed to IRE1α or ATF6 inhibitor (**Fig. 5E**). Consistent with these findings, ISRIB downregulated ATF4 levels in 7-KC exposed macrophages (**Sup. Fig. 4A**), enhanced the NET-induced DNase response and blocked the cleavage of Mertk (**Fig. 5F**) and impairment of efferocytosis (**Fig. 5G**). More importantly, similar findings were obtained in ATF4 knockdown macrophages (**Fig. 5H-J and Sup Fig. 4B**) confirming that the PERK-ATF4 branch of the ER stress signaling cascade is critical for the NET-induced impairment of DNase response in foamy atherogenic macrophages.

**Fig. 5.**
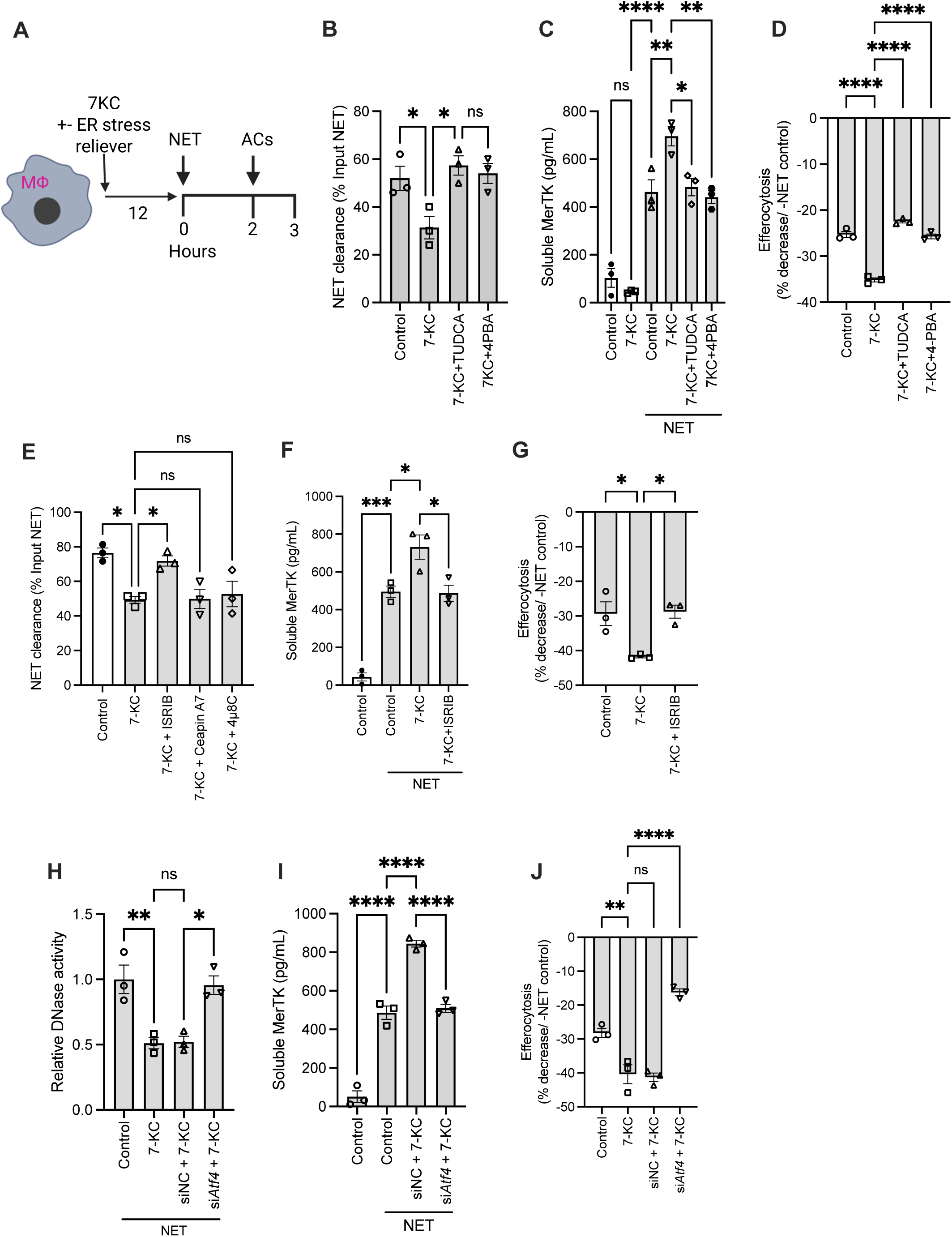
ATF4 signaling in atherogenic macrophages impairs NET-induced DNase response. **(A)** BMDMs were incubated with 7-ketocholesterol (7-KC, 15 µM) in the absence or presence of TUDCA (3 mM) or 4-PBA (0.5 μM) for 18 h. NETs (250 ng/ml) followed by fluorescently labelled apoptotic cells were added to the culture. The supernatant was used for analysis of **(B)** DNase activity as a function of NET clearance by picogreen assay, and **(C)** soluble-Mertk levels by ELISA, while the cells were used for **(D)** quantification of efferocytosis efficiency by fluorescence microscopy. n = 3 biological replicates. **(E)** BMDMs were incubated with 7-KC (15 µM) along with either vehicle, ISRIB (3 mM), Ceapin A7 (1µM), or 4µ8C (100 µM), for 18 h followed by exposure to NETs for 2 h. The NET clearance efficiency was measured by picogreen assay in the supernatant. n = 3 biological replicates. **(F)** Quantification of soluble-Mertk in the supernatants of vehicle or NET-exposed BMDMs pre-treated with 7-KC in the absence or presence of ISRIB. n = 3 biological replicates. **(G)** Quantification of efferocytosis efficiency in vehicle or NET-exposed BMDMs pretreated with 7-KC in the absence or presence of ISRIB. n = 3 biological replicates. **(H)** BMDMs transfected with negative control siRNA (siNC) or ATF4 siRNA (si*Atf4*) were incubated with vehicle or 7-KC as indicated, followed by exposure to NETs for the measurement of DNase activity, and **(I)** soluble-Mertk levels in the supernatant. n = 3 biological replicates. **(J)** Similar to (H), except that NET-exposed macrophages were incubated with apoptotic cells for the quantification of efferocytosis efficiency. n = 3 biological replicates. The data are represented as mean ± SEM. P values were calculated using ANOVA with Tukey’s multiple comparisons test (B-J). *, p < 0.05; **, p < 0.01; ***, p < 0.001.

### ISRIB boosts the NET-induced DNase response and promotes NET clearance in human and mouse atherosclerotic explant tissue

We next questioned whether these findings are relevant in the pathological context of mouse and human atherosclerosis. Towards this end, atherosclerotic aorta was harvested from *Ldlr^-/-^* mice that were fed a western diet for 16 weeks and cut into smaller fragments followed by culturing *ex vivo* as explants. The tissue was treated with either vehicle, ISRIB, or TUDCA for 4 hours prior to addition of NETs and the supernatant was collected for measurement of DNase activity and NET clearance (**Fig. 6A**). Firstly, immunoblotting of tissue extracts confirmed that treatment with both ISRIB and TUDCA decreased the ATF4 levels (**Fig. 6B**). Most importantly, both ISRIB and TUDCA led to improvement in the NET-induced DNase response (**Fig. 6C**) which was associated with accelerated clearance of NETs (**Fig. 6D**).

**Fig. 6.**
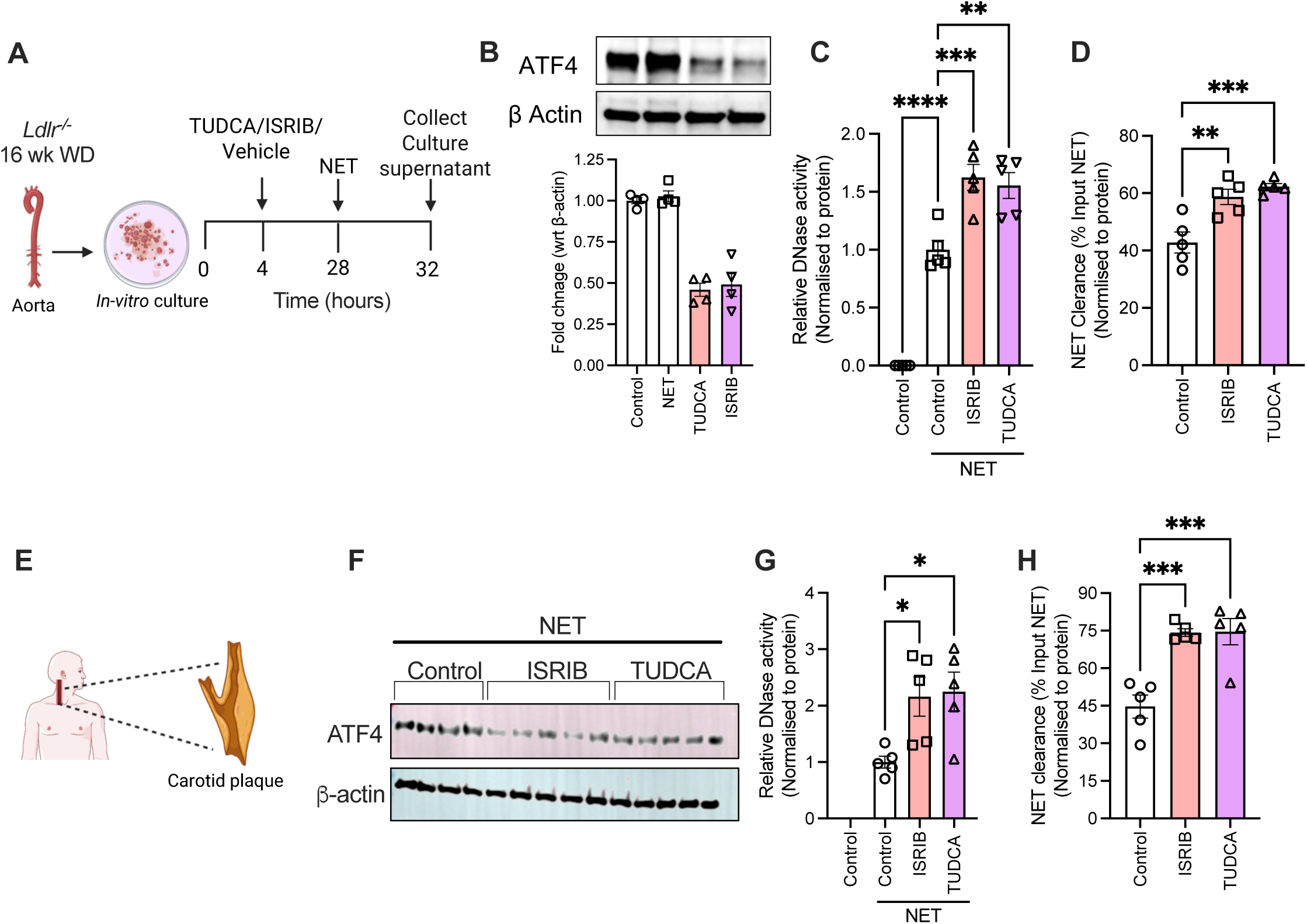
ISRIB increases NET-induced DNase response in mouse and human atherosclerotic plaques. **(A)** Aorta harvested from 16-wk WD-fed *Ldlr^-/-^* mice were cultured as explants and treated with either vehicle, TUDCA, or ISRIB (3 mM) for 18 h followed by exposure to NETs for 4 h. Tissue lysates were used for **(B)** immunoblotting of ATF4, while the culture supernatant was tested for **(C)** DNase activity, and **(D)** NET clearance efficiency. n = 4 mice per group. **(E-H)** Similar to (A-D), except that human carotid endarterectomy tissues were treated with vehicle, TUDCA, or ISRIB, followed by exposure to NETs for analysis of **(F)** tissue ATF4 levels by immunoblotting; **(G)** DNase activity, and **(H)** NET clearance efficiency in the culture supernatant. n = 5. The data are represented as mean ± SEM. P values were calculated using ANOVA with Tukey’s multiple comparisons test (B-D, G, H). *, p < 0.05; **, p < 0.01; ***, p < 0.001.

Similar to our findings above, the inhibition of ATF4 in atherosclerotic tissue explant culture from patients undergoing carotid endarterectomy by treatment with ISRIB or TUDCA (**Fig. 6E-F**) led to an increase in NET-induced DNase activity in the supernatant (**Fig. 6G**) and enhanced clearance of NETs (**Fig. 6H**). Together, these findings demonstrate that activation of the ATF4 branch of the ER stress signaling pathway mediates the impairment of the protective NET-induced DNase response in atherosclerosis.

### ISRIB improves the vascular DNase response *in vivo* and stabilizes advanced atherosclerotic plaques

Based on the *ex vivo* data above, we questioned whether inhibition of ATF4 with ISRIB could enhance DNase secretion by ER stressed lesional macrophages *in vivo* to promote the clearance of NETs in advanced atherosclerotic plaques. To address this question, *Ldlr^-/-^* mice were fed a western diet for 12 weeks to induce the generation of advanced atherosclerotic plaques and were then administered either ISRIB (1 mg/kg intraperitoneal) or vehicle for the next 4 weeks while continuing a western diet (**Fig. 7A**). Consistent with inhibition of PERK signaling, ISRIB-treated mice had lower ATF4 levels in the aorta (**Sup. Fig. 5A**). Plasma DNase activity and NET levels (**Sup. Fig. 5B-C**) and the basal aortic tissue DNase activity (**Sup. Fig. 5D**) were not different between the two groups of mice suggesting that ATF4 inhibition does not affect the total DNase levels in blood and tissues. However, aortic explants from the ISRIB-treated group demonstrated increased release of DNase in the supernatant in response to NETs (**Fig. 7B**) and increased efficiency of clearance of NETs (**Fig. 7C**) suggesting that ATF4 inhibition by ISRIB increases the NET-induced DNase response in atherosclerosis. Consistent with these *ex vivo* data, the atherosclerotic lesional NET content was significantly lower in the ISRIB-treated mice as compared with the vehicle-treated mice (**Fig. 7D**). Since ISRIB does not affect cholesterol-induced neutrophil NETosis (**Sup. Fig. 5E**), these data demonstrate that ATF4 inhibition *in vivo* increases the clearance of NETs via enhanced DNase-mediated degradation. More importantly, as predicted, the decrease in lesional NET content in the ISRIB-treated mice was associated with both higher Mertk levels in lesional macrophages (**Fig. 7E**) and increased efferocytosis efficiency (**Fig. 7F**). ISRIB treatment had no effect on the total atherosclerotic lesion area (**Fig. 7G**) which is consistent with the lack of difference in plasma cholesterol, triglycerides, and glucose between the vehicle- and ISRIB-treated mice (**Sup. Fig. 5F-I**). However, interestingly, the ISRIB-treated mice showed significant plaque remodeling as demonstrated by the decrease in plaque necrosis (**Fig. 7H**) and increased intimal collagen content (**Sup. Fig. 5J**) which is consistent with the improvement in lesional macrophage efferocytosis in these mice. Finally, the ISRIB-treated mice showed a decrease in the mRNA levels of key pro-inflammatory cytokines including *Tnf*, *Il1b*, and *Il6* (**Fig. 7I**). These data suggest that ATF4 inhibition mediated by ISRIB could promote plaque stabilization in advanced atherosclerosis via improved macrophage DNase release, NET degradation, and preservation of macrophage efferocytosis efficiency.

**Fig. 7.**
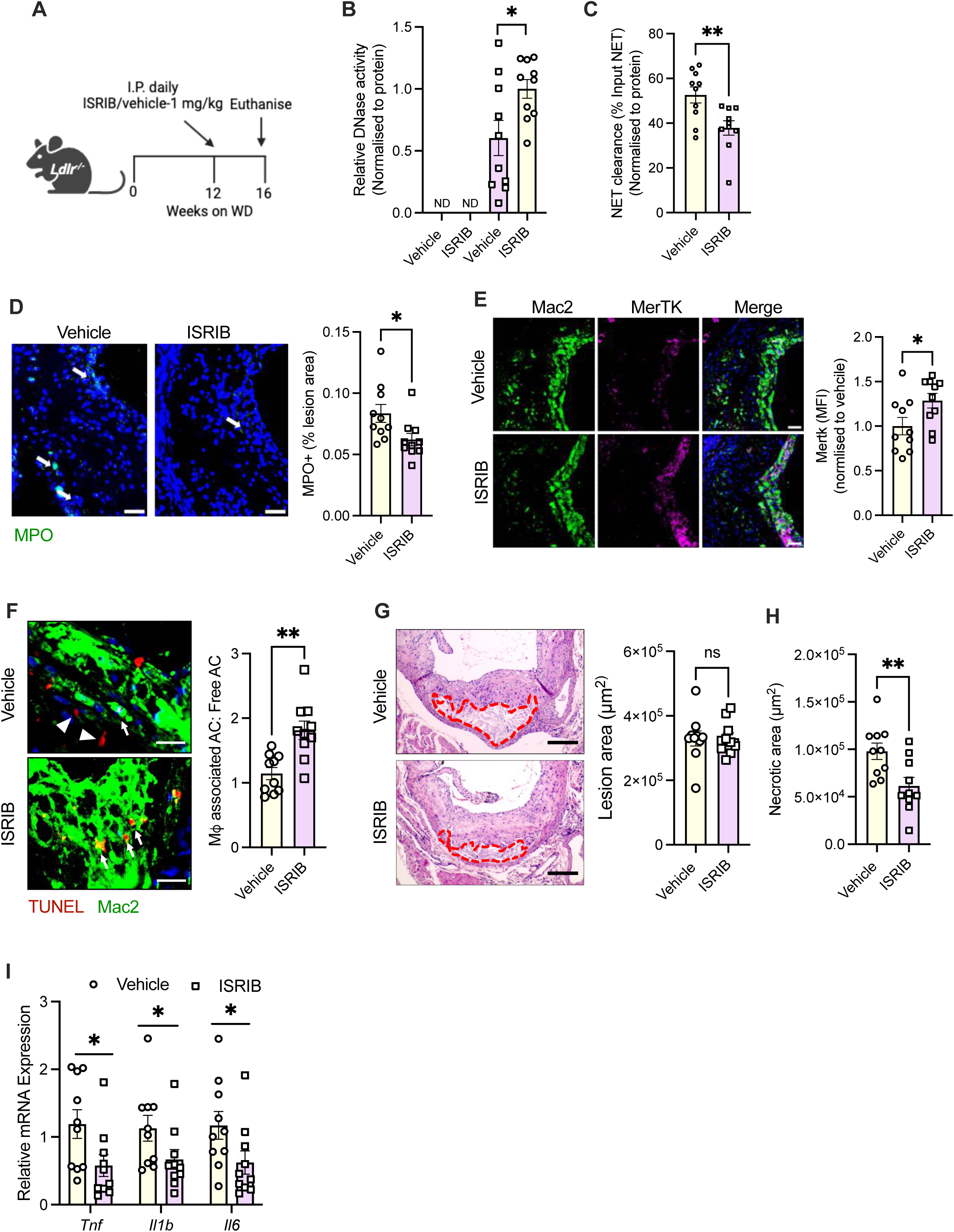
ISRIB enhances vascular DNase activity and NET clearance in murine atherosclerosis. **(A)** 10 week-old female *Ldlr^-/-^* mice were fed a WD for 16 weeks. During the final 4 weeks, one group of mice received ISRIB (1 mg/kg i.p.) daily, while the other group received vehicle. **(B)** Aortic explants from vehicle and ISRIB treated mice were exposed to NETs for measurement of DNase activity, and **(C)** NET clearance efficiency, in the supernatants. **(D)** Aortic root sections were immunostained with anti-MPO antibody for quantification of lesional NET levels, or **(E)** Mertk levels in lesional Mac2+ macrophages. **(F)** Aortic root sections were labeled with TUNEL (red) and immunostained with anti-Mac2 antibody (green) for measurement of lesional in-situ efferocytosis efficiency as a ration of macrophage-associated apoptotic cells:free-lying apoptotic cells. White arrows indicate TUNEL+ apoptotic cells associated with a macrophage. White arrowheads show free-lying TUNEL+ cells. **(G)** H&E-stained aortic root sections were used for quantification of total lesion area, and **(H)** necrotic area, in vehicle and ISRIB-treated mice. The necrotic regions are demarcated by the red dashed lines. **(I)** qPCR-based analysis of *Tnf*, *Il1b*, and *Il6*, in vascular tissues obtained from vehicle and ISRIB-treated mice. n = 10 mice per group. The data are represented as mean ± SEM. Data were tested for normal distribution using Shapiro-Wilk test. P values were calculated using unpaired t-test (B-H) and ANOVA with Tukey’s multiple comparisons test (I). *, p < 0.05; **, p < 0.01; ***, p < 0.001.

## Discussion

Our study highlights the critical role of macrophages in the clearance of NETs in inflamed tissues through the release of DNases. We demonstrate that ER stress-mediated activation of the PERK-ATF4 pathway impairs the NET-induced DNase response in lesional macrophages, resulting in defective NET clearance and their persistence in atherosclerosis. Notably, this defect can be reversed with the PERK inhibitor ISRIB. These data highlight impaired NET clearance as a key driver of NET persistence in atherosclerosis. Finally, we demonstrate that NETs promote the cleavage and shedding of Mertk on macrophages leading to decreased lesional efferocytosis. Collectively, these findings provide a mechanistic understanding of NET persistence in atherosclerosis and its role in driving plaque necrosis and instability through cellular cross talk.

NETosis is induced in the atherosclerotic plaque by multiple factors including cholesterol crystals^8^, oxidative stress^28^, and inflammatory cytokines such as CCL7^9^ and IL1β^29^. Additionally, mutations driving CHIP are associated with enhanced NETosis susceptibility^11–13^. Inhibition of NETosis by targeting neutrophil elastase^30^, PAD4^31^, MPO^32^, or the blockage of NET-associated components such histones^33^, leads to decreased atherosclerosis progression. Similarly, degradation of NETs by administration of DNases is associated with decreased atherosclerosis and plaque stabilization^18,34^. Therefore, NETosis inhibitors as well as novel synthetic DNases with high *in vivo* stability are highly sought after as therapeutic agents. However, NETs are important for host defense against pathogens and thus long-term inhibition of NETosis such as that will be required for preventing atherosclerotic complications is likely to be associated with compromised host defense and increased risk of infections as already noted, with other anti-inflammatory therapies such as anti-IL1 beta and colchicine^35,36^. In this context, our discovery of fundamental molecular mechanisms that disrupt the physiological NET-induced DNase response, such as ER stress in atherosclerotic plaque macrophages, opens new therapeutic avenues via targeting these pathways.

Lesional cells including plaque macrophages experience ER stress due to a combination of lipid accumulation, oxidative stress, and exposure to inflammatory mediators within the atherosclerotic plaque niche. ER stress perturbs macrophage function, and when excessive, triggers cell death^37^. Chemical chaperones such as 4-phenylbutyric acid (4-PBA) and TUDCA relieve ER stress in atherosclerosis and are associated with decreased plaque progression and plaque stabilization^38^. Since ER stress is a protective physiological response required for homeostatic functions including insulin secretion by pancreatic β-cells^39^, antibody production by plasma cells^40^, and dendritic cell differentiation^41^, the use of chemical chaperones such as PBA and TUDCA lead to on-target adverse effects that hinder its clinical use. Therefore, our identification of PERK-ATF4 branch of the ER stress signaling cascade as the specific trigger that impairs the NET-induced DNase response in lipid-loaded macrophages is a step change for the development of targeted therapeutic strategies. Indeed, we show that inhibition of PERK signaling using ISRIB enhances vascular DNase activity, decreases NET accumulation, and promotes plaque stabilization by decreasing lesional necrosis and enhancing collagen deposition. Given that ISRIB partially inhibits PERK signaling, it allows tempering or reshaping the aberrant ER stress responses to physiological levels and thus minimizes on-target toxicities associated with abolishment of ER stress signaling^42^. Importantly, our demonstration of conservation of these mechanistic pathways in human atherosclerosis suggests that ISRIB and similar new classes of drugs, complemented by cell- and tissue-targeted nanomedicines, provide a window of opportunity for therapeutic targeting of hyperactivated ER stress signaling cascades in chronic diseases such as atherosclerosis^43,44^.

There is extensive evidence that NETs increase plaque instability by decreasing fibrous cap thickness and increasing plaque necrosis^18^. The NET-associated decrease in fibrous cap thickness is driven by a combination of NET-induced smooth muscle apoptosis and matrix metalloprotease-mediated degradation of extracellular matrix^9^. In contrast, the mechanisms driving plaque necrosis are not well defined although an increase in NET-induced lesional cell death has been implicated. Importantly, whether NETs affect lesional efferocytosis, a major driver of atherosclerotic plaque necrosis, was unknown. Previous evidence from acute inflammatory disease settings such as sepsis^45^ suggested that NETs could impair efferocytosis. Indeed, our findings highlight that NETs decrease macrophage efferocytosis efficiency by activating ADAM17-mediated cleavage and shedding of the dominant efferocytosis receptor Mertk. While we showed previously that Mertk cleavage is a causal mechanism in driving plaque necrosis in advanced atherosclerosis^46^, the pathophysiologically relevant inducers of Mertk cleavage in the context of atherosclerosis were not known.

Recently, IL1β was described as an inducer of Mertk cleavage^14^. Adding to this repertoire, our findings establish NETs and NET-associated HMGB1 as an atherosclerotic plaque relevant inducer of Mertk cleavage. These findings raise the interesting possibility of targeting HMGB1 to preserve lesional efferocytosis efficiency. Moreover, since Mertk cleavage is a central mechanism in regulating plaque necrosis, therapeutic strategies that stabilize Mertk could be hugely beneficial in promoting plaque stabilization.

In summary, our study advances several new concepts including a) the establishment of macrophages as the major producer of DNases and as a critical determinant of the efficiency of NET clearance in tissues; b) the demonstration that activation of PERK signaling is a key pathogenic mechanism impairing the NET-induced DNase response in atherogenic macrophages; and c) the demonstration that NETs directly impair macrophage efferocytosis efficiency by cleavage of Mertk. These findings highlight several novel therapeutic strategies to enhance the clearance of lesional NETs and promote atherosclerotic plaque stabilization. Since NETs exacerbate the complications of atherosclerotic CVDs such as myocardial infarction and stroke^47,48^ as well as play a pathogenic role in several chronic inflammatory diseases, our findings could be broadly relevant for the prevention NET-induced inflammation and tissue damage in these settings.

## Materials and Methods

### Animals and Animal Maintenance

C57BL/6J mice were purchased from Charles River (UK). *Ldlr^-/-^* mice (stock # 002207) were purchased from Jackson Laboratories. *Dnase1/Dnase1l3^-/-^* mice^17^ were bred and maintained in-house. The mice were housed in IVC cages at the Biological Service Unit at Queen Mary University of London under specific pathogen-free conditions with a 12-hour light/dark cycle and had unrestricted access to food and water. All animal experimental procedures performed were approved by the Home Office, UK. Bones from Mertk^CR^ knockin mice^46^ was a kind gift from Prof Ira Tabas, Columbia University Medical Center, NY. *Tlr4^-/-^* mice^49^ were obtained from The Francis Crick Institute under an MTA from Osaka University. For the induction of atherosclerosis, *Ldlr^-/-^* mice were fed a western type diet (46% Kcal from fat and 0.2% cholesterol, from SDS, Cat # 829100) for 16 weeks.

### Neutrophil differentiation and isolation of NETs

Human HL-60 (acute promyelocytic leukemia) cells were purchased from ATCC and cultured in RPMI-1640 media supplemented with 1 mM sodium pyruvate, 10% (vol/vol) fetal bovine serum, and 100 U/mL Penicillin-Streptomycin. HL-60 Cells were seeded at ten million cells per T-75 flask and treated with 1 µM All-trans retinoic acid for 4 days to differentiate into neutrophil-like cells. For induction of NETosis, bone marrow-derived neutrophils (BMDNs) and differentiated HL-60 cells were resuspended with serum-free medium and subsequently treated with PMA (100 nM), or 7-KC (50 µg), for 4 hours.

Adherent NETs were scraped from the surface and collected, together with cells and the culture supernatant. Cells and cellular debris were separated from the supernatant using low-speed centrifugation at 450 *g* for 10 minutes at 4°C. Subsequently, the supernatant enriched with NETs was subjected to high-speed centrifugation at 18,000 *g* for 30 minutes at 4°C. The pelleted NETs were then resuspended in ice-cold PBS and stored at -80°C until further use.

### Mouse bone marrow-derived macrophages

Female or male mice aged 8-12 weeks were euthanized and their femur and tibia were collected. Bones were flushed with media and filtered through a 100 µm filter followed by centrifugation at 300 *g* for 5 minutes. RBCs were lysed by incubation with RBC lysis buffer (Sigma, R7757) for 2 minutes followed by pelleting. Cells were resuspended and cultured in DMEM high glucose supplemented with 10% fetal bovine serum, Penicillin-Streptomycin, and 20% L-929 cell culture supernatant^50^. Half the medium was replaced on day three, and cells were cultured until day seven to achieve complete macrophage differentiation.

### Aortic explant culture protocol

Mice were euthanized, and following intraventricular perfusion with PBS, the entire aorta— from the carotid artery to the iliac artery bifurcation—was dissected under a dissection microscope. The aorta was carefully cleaned to remove any excess fat from its outer layer. It was then sectioned into 1 mm fragments using a fine scalpel blade, and the fragments were randomized. The aortic explants were cultured for 4 hours in RPMI-1640 medium supplemented with 1 mM sodium pyruvate, 10% (vol/vol) fetal bovine serum, and 100 U/mL Penicillin-Streptomycin. After 4 hours, the medium was replaced with serum-free medium, and the explants were treated with or without NETs (500 ng/mL). The spent medium was collected, and residual NETs in the culture supernatant were quantified using Picogreen assay. The supernatant was concentrated 100X using a10 kDa filter followed by measurement of DNase activity by SRED. For specific experiments, the explants were either left untreated or pre-treated overnight with the ER stress relievers TUDCA (50 μg/mL) or ISRIB (0.5 μM) prior to exposure to NETs. Human carotid endarterectomy specimens from patients with symptomatic atherosclerosis were obtained after informed consent and approval of the ethics committee of Mater Misericordiae University Hospital (1/378/2372). Human plaque processing and dissection were performed as previously described^51^. Human carotid endarterectomy tissues were used for DNase assay and NET clearance as described above.

### Single radial enzyme diffusion (SRED) assay

DNase buffer (20 mM Tris-HCl, pH 7.8, 10 mM MnCl2, 2 mM CaCl2) was prepared and mixed with salmon testes DNA (55 µg/mL) and 2X SYBR Safe. The mixture was incubated at 50°C for 10 minutes, followed by the addition of an equal volume of 2% agarose dissolved in nuclease-free water. The resulting solution was poured into plastic trays and allowed to solidify at room temperature. Wells approximately 0.1 mm in size were created in the gel using a 20 µL pipette tip. Subsequently, 2 µL of plasma, 5 µL of 100X concentrated cell culture supernatant and peritoneal fluid, or 5 µL of tissue extract were loaded into the wells. Tissue extracts for DNase activity were prepared by homogenizing tissues using bead homogenization (VP 43 SOP) in Tris-HCl, pH 7.8. The crude extracts were centrifuged at 20,000 *g* for 10 minutes at 4°C, and the supernatants were collected. The gels were incubated for 6 hours at 37°C in a humidified chamber. After incubation, images of the gels were captured using the Azure 400 gel documentation system. DNA degradation was quantified using Fiji software by comparing the samples against a standard curve generated from known concentrations of purified DNase1 run on the same gel.

### Denaturing polyacrylamide gel electrophoresis zymography (DPZ)

To measure individual DNase1 and DNase1L3 activity, 10% (v/v) resolving gels were made by mixing with DNA extracted from salmon testes (200 μg/mL). For DNase1 activity assessment, 0.5 µL and 5 µL of 100X concentrated cell culture supernatant and peritoneal exudate were mixed with 12 µL of nuclease-free water and 5 µL of SDS gel-loading buffer. The solution was heated for 5 minutes and then loaded onto the gels. After electrophoresis, gels were washed twice with 10 mM Tris/HCl pH 7.8 at 50°C for 30 minutes to remove remnant SDS. Gels were transferred into refolding buffer (10 mM Tris/HCl pH 7.8, 3 mM CaCl2, 3 mM MgCl2) and incubated at 37°C overnight.

For DNase1L3 detection, 2 µL of serum was used, and samples were mixed with 1 µL of beta-mercaptoethanol (BME) before loading onto the gel. Electrophoresis and subsequent washing steps were carried out as described above, and the gel was incubated for 48 hours at 37°C in refolding buffer-1 (10 mM Tris/HCl pH 7.8, 1 mM BME), followed by another 48- hour incubation at 37°C in refolding buffer-2 (refolding buffer 1 supplemented with 3 mmol/L CaCl2 and 3 mmol/L MnCl2). Images were captured using a Gel Doc system (FluorChem E), and quantification was conducted using Fiji software.

### Quantification of NETs by MPO-DNA ELISA

To quantify NETs, polyclonal anti-myeloperoxidase antibodies were coated onto 96-well ELISA plates using a carbonate-bicarbonate buffer at a dilution of 1:1000 for 12 hours at 4°C. Following a 1-hour blocking step at room temperature (RT) with 3% bovine serum albumin (BSA), the wells were incubated for 12 hours at 4°C with 50 µL of plasma, 200 µL of peritoneal exudate, and 200 µL of cell culture supernatant. After three washes with PBS containing 0.1% Tween 20 (PBST), the DNA content was quantified using the Quant-iT PicoGreen dsDNA assay kit, following the manufacturer’s instructions. For positive and negative controls, appropriate wells were incubated with *in vitro* generated NETs and genomic DNA, respectively.

### Soluble MerTK ELISA

Soluble MerTK levels were measured using a Mouse Mer DuoSet ELISA kit (R&D Systems) following the manufacturer’s instructions. ELISA plates were coated with a capture antibody overnight, blocked with 1% BSA, and incubated with samples or standards for 2 hours at room temperature. After adding the detection antibody conjugated to HRP-streptavidin, the TMB substrate was applied, and the reaction was stopped. Absorbance was measured at 450 nm.

### *In vitro* efferocytosis assay

HL-60 cells were labelled with CellVue Claret according to the manufacturer’s protocol. Briefly, cells were washed with PBS and then resuspended in 2 mL of Diluent C along with 4 µL of the dye. CellVue Claret-labeled cells were exposed to UV-C radiation (254 nm) for 7 minutes, equivalent to 5 kJ/m^2^, followed by a 1-hour incubation at 37°C to induce apoptosis. BMDMs were plated at a density of 0.5 million cells per well in a 12-well plate. Apoptotic HL60 cells were added at a ratio of 1:1 ACs to Macrophages and incubated for 1 hour. After the incubation period, unengulfed ACs were washed away, and images were captured using a fluorescence microscope (EVOS FL) at 20X magnification. FIJI (ImageJ) was used to quantify efferocytosis efficiency by calculating the percent macrophages that have engulfed an AC.

### Quantification of efficiency of NET clearance

For the NET clearance assay, BMDMs were incubated with NETs (250ng/mL) in serum-free media. At the specified time points for each experiment, the amount of remaining NETs in the supernatant was quantified using the Quant-iT PicoGreen dsDNA assay kit, following the manufacturer’s protocol. The percentage of NETs remaining was calculated relative to the initial amount of NETs added.

### RNA Isolation, cDNA preparation and qPCR

Total RNA was extracted from cells using the RNeasy Mini Kit according to the manufacturer’s instructions. For total RNA extraction from the aorta, the heart and entire aorta were perfused, placed on a silicone plate, and pinned. Excess fat was removed, and the aortic arch was isolated and transferred into Precellys tubes containing 500 µL of Trizol. The tissue was homogenized using a Precellys 24 Tissue Homogenizer at 6500 rpm for 30 seconds across 20 cycles. Total RNA was then isolated using the chloroform-isopropanol method^52^. For mRNA expression analysis, 1 µg of total RNA was reverse-transcribed into cDNA using the PrimeScriptTM 1st Strand cDNA Synthesis Kit (TaKaRa). The cDNA was mixed with KAPA SYBR Green master mix, and qPCR was performed using the Roche Lightcycler 480. The results were obtained as cycle threshold (Ct) values, which were used to calculate 2-ΔΔCt and determine fold changes, providing relative mRNA expression levels normalized to an endogenous housekeeping gene.

### Bone marrow transplantation

Ten-week-old female *C57BL/6* or *Ldlr^-/-^* recipient mice underwent lethal X-ray irradiation, administered in two equal doses of 4 Gy, separated by a 4-hour interval. After the second radiation dose, 5 million donor bone marrow cells were delivered via tail vein injection. Following a 6-week period for bone marrow reconstitution, the mice were used for subsequent experiments.

### *In vivo* DNase response and efferocytosis in the peritoneal cavity

To quantify the NET-induced DNase response in the peritoneal cavity, appropriate groups of mice were administered either NETs (1mg) or a vehicle control (PBS) intraperitoneally. 3 hours later, the mice were euthanized, and peritoneal lavage was performed with 5 ml of cold PBS. The peritoneal lavage fluid was centrifuged at 18,000 *g* for 10 minutes and concentrated 100X for analysis of DNase activity using SRED and DPZ.

For analysis of efferocytosis efficiency, appropriate groups of mice were injected 10^6^ CellVue Claret-labeled apoptotic cells in 300μL of PBS into the peritoneal cavity of mice that were previously exposed to NETs or vehicle for 2 hours. One hour after injection of the apoptotic cells, the mice were euthanized, and peritoneal lavage performed. The cells in the peritoneal lavage were immunostained with fluorescently-labeled anti-F4/80 antibody. Efferocytosis was analysed by identifying the F4-80+ CellVue Claret+ population by flow cytometry.

### Adoptive transfer of macrophages into peritoneal cavity

Age-matched WT and *Dnase1/Dnase1l3^-/-^* mice resident peritoneal macrophages were isolated using the macrophage enrichment kit (Miltenyi Biotech), and 0.5×10^6^ macrophages were injected intraperitoneally into the recipient mice. 16 hours post adoptive cellular transfer, the mice were injected with NETs and apoptotic cells as described above to quantify the DNase response and macrophage efferocytosis efficiency.

### Atherosclerotic plaque morphometry, immunofluorescence, and *in situ* efferocytosis assay

At the end of experiment, the mice were euthanized, blood collected, and tissues perfused with 1X PBS via intraventricular injection. The heart, along with the intact aortic root, was fixed in 10% formalin, embedded in paraffin, and sectioned into 8 µm slices using a microtome. Fifty consecutive sections were obtained from the initial appearance of the aortic valve. Hematoxylin and eosin (H&E) staining of six equally spaced sections per mouse covering the entire aortic root were analyzed for quantification of total lesion area and necrotic area. Additionally, sections were stained with Mason’s trichrome dye (Sigma-Aldrich) following the manufacturer’s protocol to quantify lesional collagen content.

For immunostaining of FFPE (Formalin Fixed Paraffin Embedded) specimens, deparaffinization was conducted using histoclear, followed by rehydration with graded ethanol and final rinses with 1X PBS. Antigen retrieval was achieved incubating sections in sodium citrate buffer (10 mmol/L Sodium citrate, 0.05% Tween-20, pH 6.0) at 95°C for 45 minutes. After blocking with 3% FBS in 1X PBS for 30 minutes, the sections were incubated overnight at 4°C with appropriate primary antibody. Following washes, the sections were incubated with appropriate fluorescently labelled secondary antibody for 1 hour at room temperature and counterstained with DAPI.

For *in situ* efferocytosis assay, TUNEL staining was performed according to the manufacturer’s instructions. Following TUNEL labelling, the sections were immunostained with anti-Mac2 antibody as described above and counterstained with DAPI. Images were captured using a fluorescence microscope (EVOS FL), and the *in situ* efferocytosis efficiency was determined as the ratio of TUNEL+ apoptotic cells that are associated with a Mac2+ macrophage (associated) or lying free (Associated:Free). The analysis of all atherosclerotic lesions was conducted in a blinded manner, with one individual executing the experiment and another person performing the analysis.

### siRNA transfection

Lipofectamine RNAimax transfection reagent was used for siRNA transfection following the manufacturer’s protocol. Briefly, 2×10^6^ HL-60 cells were seeded in 6-well plates and treated with ATRA for 4 days prior to proceeding with knockdown. Similarly, 0.5×10^6^ BMDMs were seeded in 12-well plates and incubated overnight. To generate siRNA-lipofectamine complexes, Lipofectamine RNAimax reagent was diluted in Opti-MEM and combined with 10 pmol of siRNA targeting ATF4 or HMGB1 in an equal volume of Opti-MEM. This mixture was incubated for five minutes at room temperature and then added to cells. After 12-hour culture in Opti-MEM, cells were transferred to DMEM supplemented with 10% (vol/vol) FBS and 100 U/mL penicillin-streptomycin for 24 hours before proceeding with further experiments.

### Immunofluorescence staining of NETs

24-well plates were pre-coated with poly-L-lysine for 1 hour prior to seeding with 0.5×10^6^ ATRA-differentiated HL60 cells followed by incubation overnight. The culture media was replaced with fresh serum free media and treated with 100 nM PMA for 4 hours followed by fixation with 4% paraformaldehyde.

For immunostaining of NETs in the peritoneal exudate, 500 µL of the exudate was incubated with poly-L-lysine coated dishes at 4°C. Samples were then blocked with 3% FBS in 1X-PBS, followed by incubation with primary antibody against CitH3, MPO, or HMGB1 and subsequent incubation with fluorophore-conjugated secondary antibody. Nuclei were counterstained using DAPI. Images were captured using a fluorescence microscope and analyzed with FIJI software.

### Western blotting

Cells were lysed in 2X laemmeli buffer with 1% v/v Dithiothreitol (DTT) and heated at 95°C for 10 minutes. Samples were then either stored at -20°C or used immediately. Samples were loaded onto a 10% polyacrylamide gel along with a pre-stained ladder and electrophoresis was performed. Proteins were transferred to a PVDF Membrane via wet transfer at 90 volts for 90 minutes. The membrane was blocked with 5% milk or 5% BSA for 1 hour at room temperature, followed by brief washing. Primary antibodies were then incubated overnight at 4°C and after washes, the membrane was incubated with the appropriate secondary horseradish peroxidase (HRP)-conjugated antibody for 1 hour at room temperature. Following further washing, 1 mL of western blotting HRP substrate was used for chemiluminescent imaging using an Azure 400 gel documentation system.

### Mass-Spectrometric Analysis

NETs were digest by using sequencing grade trypsin (1:10 ratio,Trypsin:protein) for 16-18 hours at 37^0^C . Prior to digestion, sample was reduced with 25 mM DTT for 30 min at 56°C and the cysteines were blocked by 55 mM IAA at room temperature for 15–20 min. Digested peptides were reconstituted in 5µl of LC-MS grade water containing 0.1% formic acid and run on a quadrupole-TOF hybrid mass spectrometer (TripleTOF6600, Sciex, USA) coupled to a nano-LC system (Eksigent NanoLC-400). Peptides were loaded onto a reverse phase peptide ChromoLC trap (200 mm 0.5 mm) column and separated using a C18 column (75 mm 15 cm, Eksigent). The samples were run using buffers A (99.9% LC-MS water + 0.1% formic acid) and B (99.9% acetonitrile + 0.1% formic acid) as the mobile phases. At a constant flow rate of 250 nL min-1, the gradient is composed of 95% buffer A for 2 minutes, 90% buffer A for 8 minutes, 20% buffer A for 42 minutes, and then 95% buffer A for 16 minutes. Data were acquired with a Nano Spray source installed in the TripleTOF 6600 System using a nebulizing gas of 20 psi, a curtain gas of 25 psi, an ion spray voltage of 2000 V and a heater interface temperature of 75 ^0^C. Data-dependent acquisition (DDA) mode was setup with a TOF/MS survey scan (350–1600 m/z) with an accumulation time of 250 ms. A maximum of ten precursor ions per cycle were chosen for fragmentation, with each MS/MS spectrum (200–1800 m/z) being accumulated for 70 ms, for a total cycle time of roughly 2.05 seconds. Parent ions were chosen for MS/MS fragmentation if their abundance was greater than 150 cps and their charge state ranged from +2 to +5. The protein pilot v5.0 was used to analyse the .wiff files produced by the Triple-TOF 6600, which include both MS and MS/MS spectra, to identify the desired protein.

### Statistical analysis

All data are presented as mean ± SEM. The number of biological replicates or the number of mice used in each experiment is indicated in the figure legend. Data that passed the Shapiro-Wilk test for normal distribution were analyzed using a parametric test. Non-parametric tests were used for data that did not show normal distribution. The specific test used for each analysis is indicated in the figure legend.

## Acknowledgements

This study was supported by funding from the British Heart Foundation (PG/22/11226), UKRI-BBSRC (BB/Y513143/1), and Barts Charity (MGU0459), awarded to M.S. T.V was supported by a University College Dublin Ad Astra PhD studentship. C.G and E.B are supported by Science Foundation Ireland Awards (21/US/3751) and by the UCD Foundation. E.B and M.O’D are supported by a UCD School of Medicine COINTREAU award. We sincerely acknowledge Dr Santosh R. Sukka and Prof Ira Tabas, Columbia University Medical Center, NY, for providing bones from Mertk-cleavage resistant mice. *Tlr4^-/-^* mice were kindly provided by Prof Shizuo Akira, Osaka University.

## Supplementary Figure Legends

**Sup. Fig. 1.**
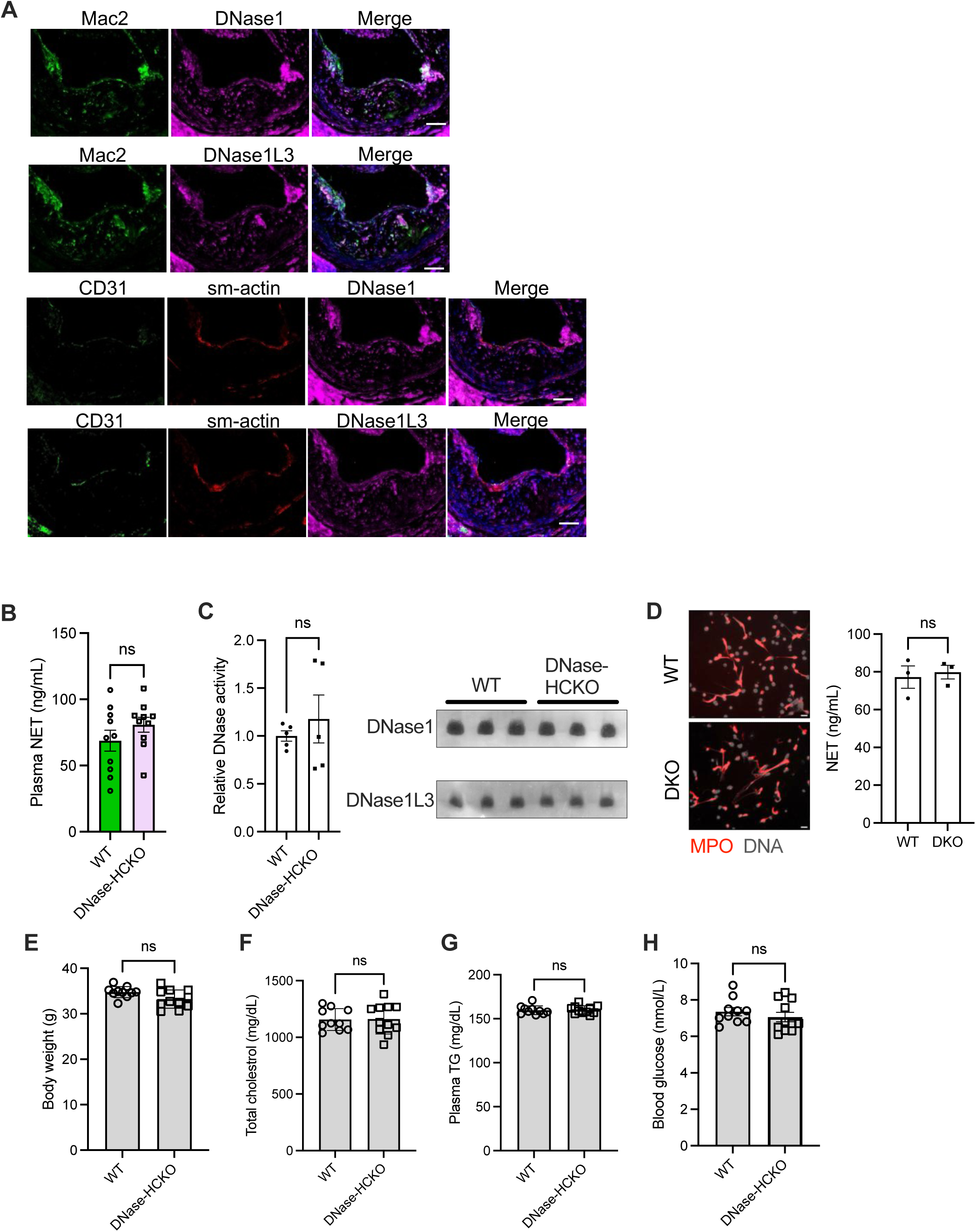
**(A)** Representative fluorescence microscopy images of aortic root sections from 16 wk WD-fed *Ldlr^-/-^* mice immunostained with anti-Mac2, anti-sm-actin, anti-CD31, anti-DNase1, and anti-DNase1L3 antibody. **(B)** Measurement of plasma DNase activity in 16 wk WD-fed WT and DNase-HCKO *Ldlr^-/-^* mice. **(C)** Analysis of relative levels of plasma DNase1and DNase1L3 by DPZ in 16 wk WD-fed WT and DNase-HCKO *Ldlr^-/-^* mice. **(D)** Bone marrow neutrophils isolated from WT and DKO mice were exposed to PMA to induce NETosis. The left panel shows representative fluorescence microscopy images after labeling cells with anti-MPO antibody and SYTOX green. Right panel shows quantification of NET levels by MPO-DNA ELISA. **(E)** Body weight of WT and DNase-HCKO *Ldlr^-/-^* mice at 16 wks of WD feeding. **(F)** Quantification of total cholesterol, and **(G)** triglycerides in plasma of 16 wk WD-fed WT and DNase-HCKO *Ldlr^-/-^* mice. **(H)** Analysis of blood glucose levels in 16 wk WD-fed WT and DNase-HCKO *Ldlr^-/-^* mice. n = 10 mice per group. The data are represented as mean ± SEM. Data were tested for normal distribution using Shapiro-Wilk test. P values were calculated using unpaired t-test (B, D-H).

**Sup. Fig. 2.**
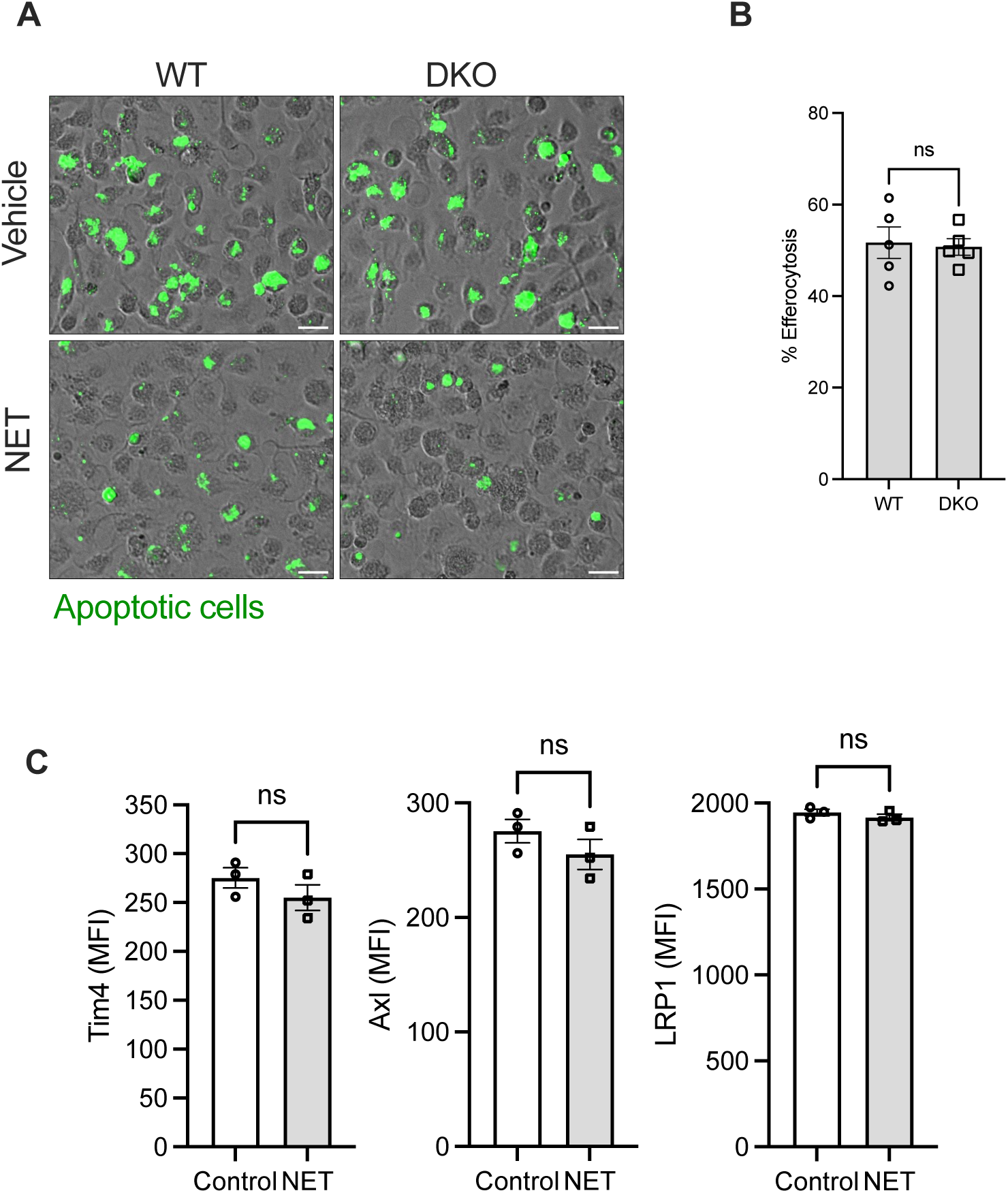
**(A)** WT and DKO BMDMs were exposed to vehicle or NETs for 2 h followed by incubation with fluorescently labelled apoptotic cells for quantification of efferocytosis efficiency. Representative fluorescence microscopy images are shown. **(B)** The bar graph shows efferocytosis quantification of WT and DKO BMDMs under homeostatic conditions. **(C)** The bar graphs represent flow cytometric quantification of cell surface levels of Tim4, Axl, and LRP1, in macrophages exposed to vehicle or NETs.

**Sup. Fig. 3.**
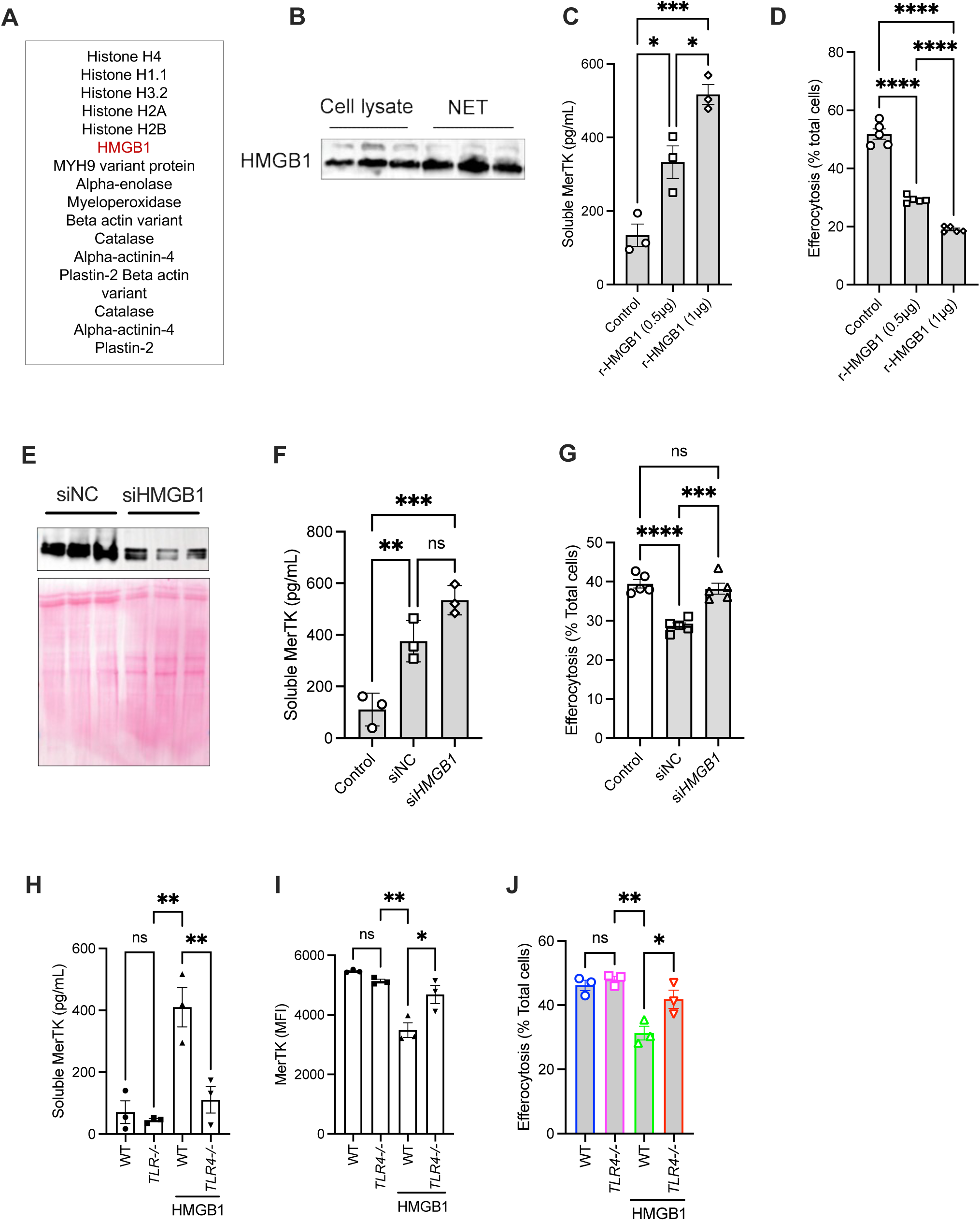
**(A)** The table shows a list of NET-associated proteins detected by mass spectrometry. **(B)** Immunoblotting for HMGB1 in ATRA-differentiated HL60 cell lysates and NETs. **(C)** BMDMs were incubated with recombinant HMGB1 for 2 h followed by measurement of soluble-Mertk in the supernatant. n = 3 biological replicates. **(D)** As above, except that HMGB1 exposed macrophages were incubated with apoptotic cells for quantification of efferocytosis efficiency. n = 4 biological replicates. **(E)** Differentiated HL-60 cells transfected with siNC or siHMGB1 were cultured for 24 h before PMA-induced NETosis. NETs were analyzed by western blot for HMGB1. Bottom panel shows Ponceau S-stained membrane. **(F-G)** Soluble Mertk levels and efferocytosis efficiency was analyzed in BMDMs incubated with NETs derived from HL60 cells transfected with siNC or siHMGB1. **(H-J)** Soluble-Mertk levels, cell Mertk levels, and efferocytosis efficiency was quantified in WT and *Tlr4^-/-^* BMDMs exposed to recombinant HMGB1(0.5 µg) for 2 h. The data are represented as mean ± SEM. P values were calculated using ANOVA with Tukey’s multiple comparisons test (C, D, F-J). *, p < 0.05; **, p < 0.01; ***, p < 0.001.

**Sup. Fig. 4.**
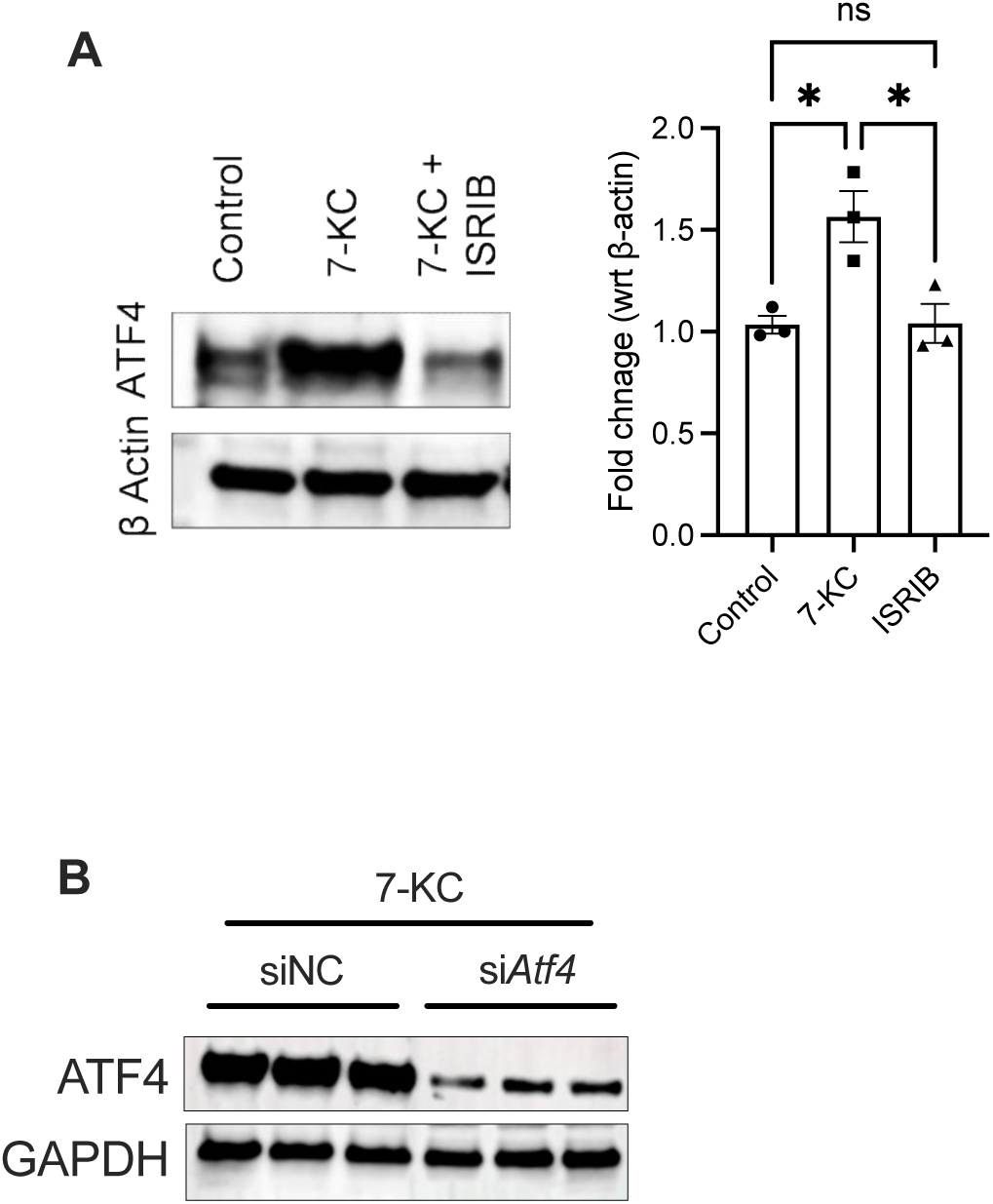
**(A)** Immunobloting for ATF4 in lysates obtained from BMDMs treated with 7-KC in the absence or presence of ISRIB. **(B)** Immunobloting for ATF4 in siNC or si*Atf4* transfected BMDMs treated with 7-KC. n = 3 biological replicates. The data are represented as mean ± SEM. P values were calculated using ANOVA with Tukey’s multiple comparisons test (C, D, F, G). *, p < 0.05; **, p < 0.01; ***, p < 0.001.

**Sup. Fig. 5.**
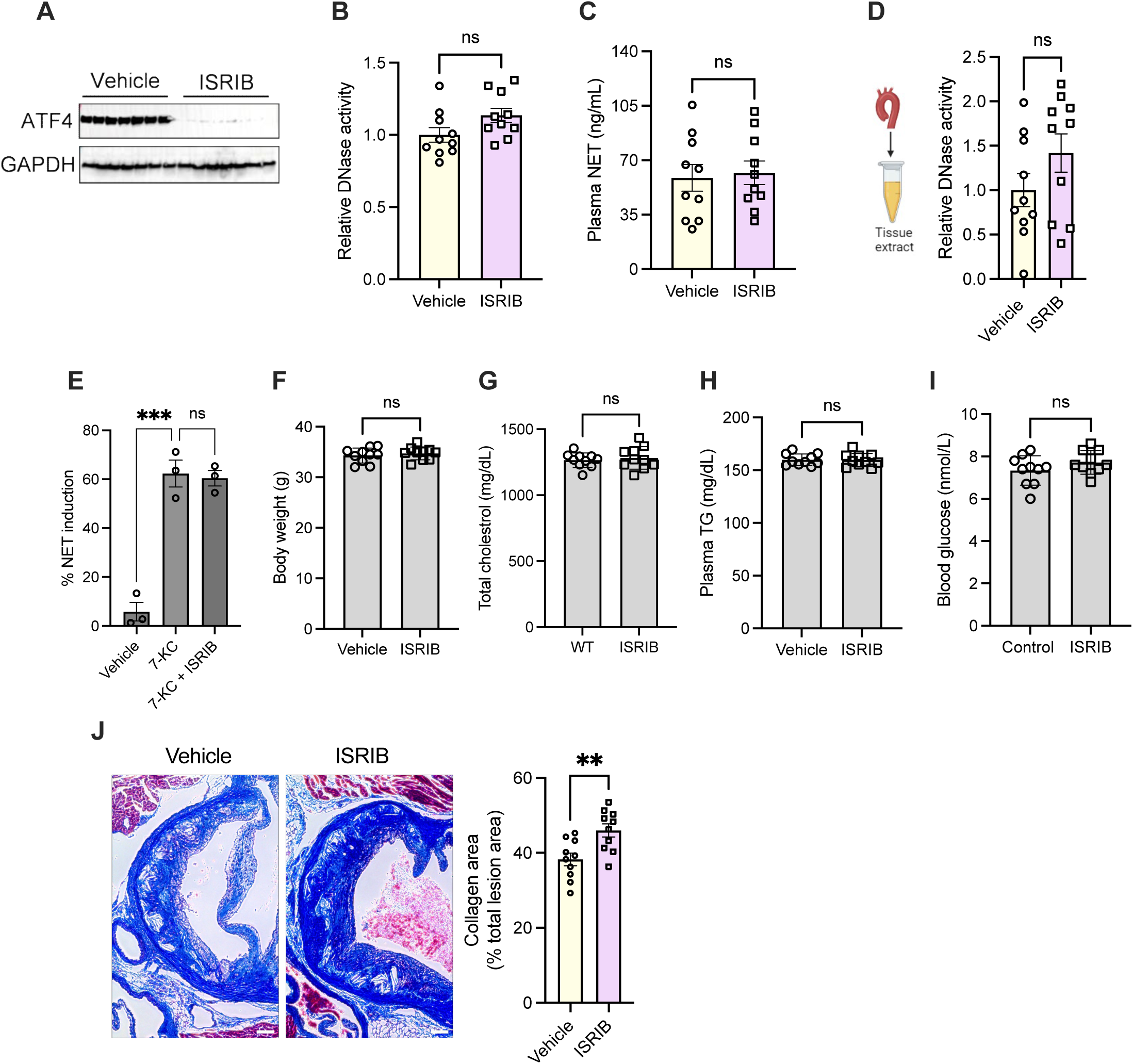
*Ldlr^-/-^* mice fed a WD for 16 weeks were treated with either vehicle or ISRIB for the last 4 weeks. The following parameters were quantified: **(A)** ATF4 levels in aortic tissue by western blotting; **(B)** plasma DNase activity; **(C)** Plasma level of NETs; **(D)** basal DNase activity in the aortic tissue; **(E)** Quantification of 7-KC-induced NETosis in vehicle and ISRIB-treated neutrophils; **(F)** body weight; **(G)** plasma total cholesterol; **(H)** plasma triglycerides; **(I)** blood glucose; and **(J)** Quantification of lesional collagen content by Mason’s trichrome staining in aortic root sections of vehicle and ISRIB treated *Ldlr^-/-^* mice. n = 10 mice per group. The data are represented as mean ± SEM. Data were tested for normal distribution using Shapiro-Wilk test. P values were calculated using unpaired t-test (A, B, D-J).

